# Nucleoid-Associated Proteins Undergo Liquid-Liquid Phase Separation with DNA into Multiphasic Condensates Resembling Bacterial Nucleoids

**DOI:** 10.1101/2022.06.23.497280

**Authors:** Archit Gupta, Ashish Joshi, Kanika Arora, Samrat Mukhopadhyay, Purnananda Guptasarma

**Author notes:** Contributed equally (Archit Gupta, Ashish Joshi).

## Abstract

Liquid-liquid phase separation offers unique spatiotemporal control over myriad complex intracellular biochemical processes through compartmentalization of biomolecules into highly dynamic, liquid-like condensates known as membrane-less organelles. The bacterial nucleoid is thought to be one such phase-separated condensate; however, its formation, regulation, and biophysical characteristics are poorly understood. Our super-resolution imaging data suggests that nucleoids are dynamic assemblages of sub-micron-sized liquid-like droplets. We demonstrate that non-sequence-specific <u>N</u>ucleoid-<u>A</u>ssociated <u>P</u>roteins (NAPs) such as HU-A, HU-B and Dps, accrete nucleic acids and spontaneously condense with them into liquid-like, multicomponent, multiphasic, heterotypic phase-separated droplets. Upon mixing of HU-B, DNA and Dps, HU-B-enriched droplets are seen to contain demixed Dps-enriched droplets. Our findings indicate scope for the possible existence of multiphasic liquid-like compartments within nucleoids, providing insights into bacterial growth phase-dependent variations in the levels of different NAPs.

## Introduction

Biomolecular condensation of proteins and nucleic acids through liquid-liquid phase separation (LLPS) is thought to exert spatiotemporal control over complex intracellular biochemical processes and biological phenomena.^1–7^ LLPS is proposed to drive the formation of non-stoichiometric, highly dynamic, liquid-like, mesoscopic, subcellular, membrane-less compartments, both within the cytoplasm and in the nucleus^8,9^ of eukaryotic cells, e.g., in respect of stress granule formation^10^, nucleolar organization,^11^ replication, transcription, and translation regulation.^12^ In prokaryotic cells too, recent exciting developments have established that critical cellular processes are governed by LLPS, e.g., subcellular organization, chromosomal segregation, transcriptional and translation regulation,^13,14^ cell division,^15^ and DNA protection.^16^ The self-assembly of DNA in mitochondria (which are thought to be eukaryotic cellular organelles of prokaryotic origin) is also LLPS-dependent. ^17^

Bacterial nucleoids contain genomic DNA (~ 80%), nucleoid-associated proteins (~ 10%), and RNA (~ 10%).^18^ Their leitmotif is their ability to persist through successive rounds of replication and transcription, spanning a virtually infinite series of cell generations, while remaining in an extreme state of compaction. This compaction facilitates the packing of a few millimeters length of double-stranded genomic DNA into a dynamic, functionally-active nucleoid inside a cell that is only about one micrometer long.^19^ The >1000-fold compaction of DNA appears not to owe to supercoiling, bending or looping of DNA alone, but also to macromolecular crowding,^20^ associated with Liquid-Liquid phase sepaation.^21^ Measurements of optical refractivity,^22^ and physical compressibility,^23^ underlie the belief that bacterial nucleoids exist in a constant state of liquid-like phase separation.

In this work, our aim was to identify key Nucleoid Associated Proteins (NAP) that could be responsible for causing DNA to enter into a phase separated state. We argued from first principles that an NAPs helping in creating and/or maintaining a state of phase separation involving the entire bacterial chromosome would have to be (a) highly conserved across bacteria, (b) highly abundant, (c) non-sequence-specific in its binding of nucleic acids, and (d) characterized by the presence of some structural disorder, since disorder is thought to play a role in phase separation.^5,9,24^ Further, (e) we argued that two types of NAPs could be required for packaging DNA into a phase separated state within the nucleoid: (i) an NAP with a basic isoelectric point, which would be amenable to becoming completely buried by DNA (e.g., within the nucleoid’s interior), and (ii) an NAP with an acidic isoelectric point, which would possess an overall negative charge and resist complete burial by DNA (e.g., at nucleoid-cytoplasm interfaces).

Interestingly, application of the above criteria to the dozen or so NAPs present in the model bacterium, *Escherichia coli*, reveals that there are two NAPs satisfying the fifth criterion which fulfill the first four criteria. The basic NAP is HU; a histone-like protein first isolated from *E. coli* strain U93,^25–27^ which is constituted of two highly homologous isoforms, HU-A and HU-B, and which exists as homodimers or heterodimers of these isoforms. The acidic NAP is Dps; a DNA binding protein from starved cells, which happens to be a dodecameric homolog of ferritin.^28^ The structures of both HU and Dps contain intrinsically disordered regions.^25,28^The sequences and structures of both HU and Dps are highly conserved across bacteria.^26,29^ HU and Dps are both highly abundant, being amongst the most abundant of all bacterial proteins.^30^ HU and Dps bind to multiple physical and chemical forms of nucleic acids,^31,32^ with no known sequence-specificity.^33,34^ Dps is also a component of HU’s genetic stress response regulon^32^ and both of these are regulated by another NAP called Fis.^35^ Based on entirely *a priori* considerations, therefore, HU and Dps would appear to be the most ideal of all bacterial protein candidates to drive the formation and maintenance of a phase-separated state involving the bacterial chromosome.

We examined whether HU is associated with LLPS-like behaviour in bacterial nucleoids *in* vivo, and also whether HU and Dps engage in LLPS with DNA to form biomolecular condensates *in vitro*. Below, we present results of investigations demonstrating (a) that dynamic submicron-sized bead-like domains (imaged using genetic fusions of fluorescent proteins and HU), exist within *E. coli* nucleoids, and (b) that both individually and collectively, HU-A, HU-B, and Dps, cause the condensation of nucleic acids into phase-separated droplets, under conditions mimicking the temperature, pH, ionic strength, and concentrations of these NAPs, and nucleic acids, in *E. coli* cells. The three NAPs form complex coacervates that appear to be model *ex vivo* simulacrums of bacterial nucleoids.

## Results

### Bacterial nucleoids contain sub-micron-sized liquid-like droplets

To directly observe morphological details of *E. coli* nucleoids, we performed microscopy experiments involving live cells in which structured illumination microscopy (SIM) was used to image nucleoids inside *E. coli* cells expressing the fluorescent DNA-binding HU construct, RFP-HU-A, which we have created and used previously.^36–38^ The upper panel of Fig. 1**a** shows a DIC image of a field of *E. coli* cells containing some cells that exist as filaments owing to incomplete separation after cell division (a known outcome of overexpression of either HU, or RFP-HU-A).^37^ The lower panel of Fig. 1**a** shows a magnified DIC image of one such filament, emphasizing that such filaments are linear. Fig. 1**b** shows a super-resolution image of a linear filament of four conjoined cells, joined to a fifth (terminal) cell, placed at an angle (and apparently on the verge of separating away). All five cells contain RFP-HU-A-labelled fluorescent nucleoids. Each nucleoid consists of what appear to be multiple (sub-micron-sized) bead-like droplet entities. Each nucleoid appears to have a different shape. Fig. 1**c** shows a still image representing yet another such field of cells, showing a linear filament of conjoined *E. coli* cells. We monitored one such filament continuously for several tens of seconds, using super-resolution videography (Supplementary Information Video 1). In the video, sub-micron-sized bead-like droplet entities are indeed observed to undergo continuous remodeling over timescales measurable in seconds, inside each nucleoid. The remodeling is suggestive of fusion and separation of bead-like entities, partitioning and re-partitioning the nucleoid into new shapes inside growing cells. Overall, the data appears to be reminiscent of liquid droplets that display the tendency to fuse and also separate (or drip), except that the sizes here are ‘sub-micron’ whereas liquid droplets formed by proteins *in vitro* typically have diameters measured in microns. Given that sub-micron-sized phase-separated droplets have previously been reported by other authors,^39^ in particular, in respect of carboxysomes within bacteria,^40^ and given the possibility that micron-sized (or larger) droplets could be formed by the same entities *in vitro*, we proceeded to examine whether recombinantly-expressed forms of HU cause the formation, and maintenance, of a phase separated state of DNA.

**Figure 1.**
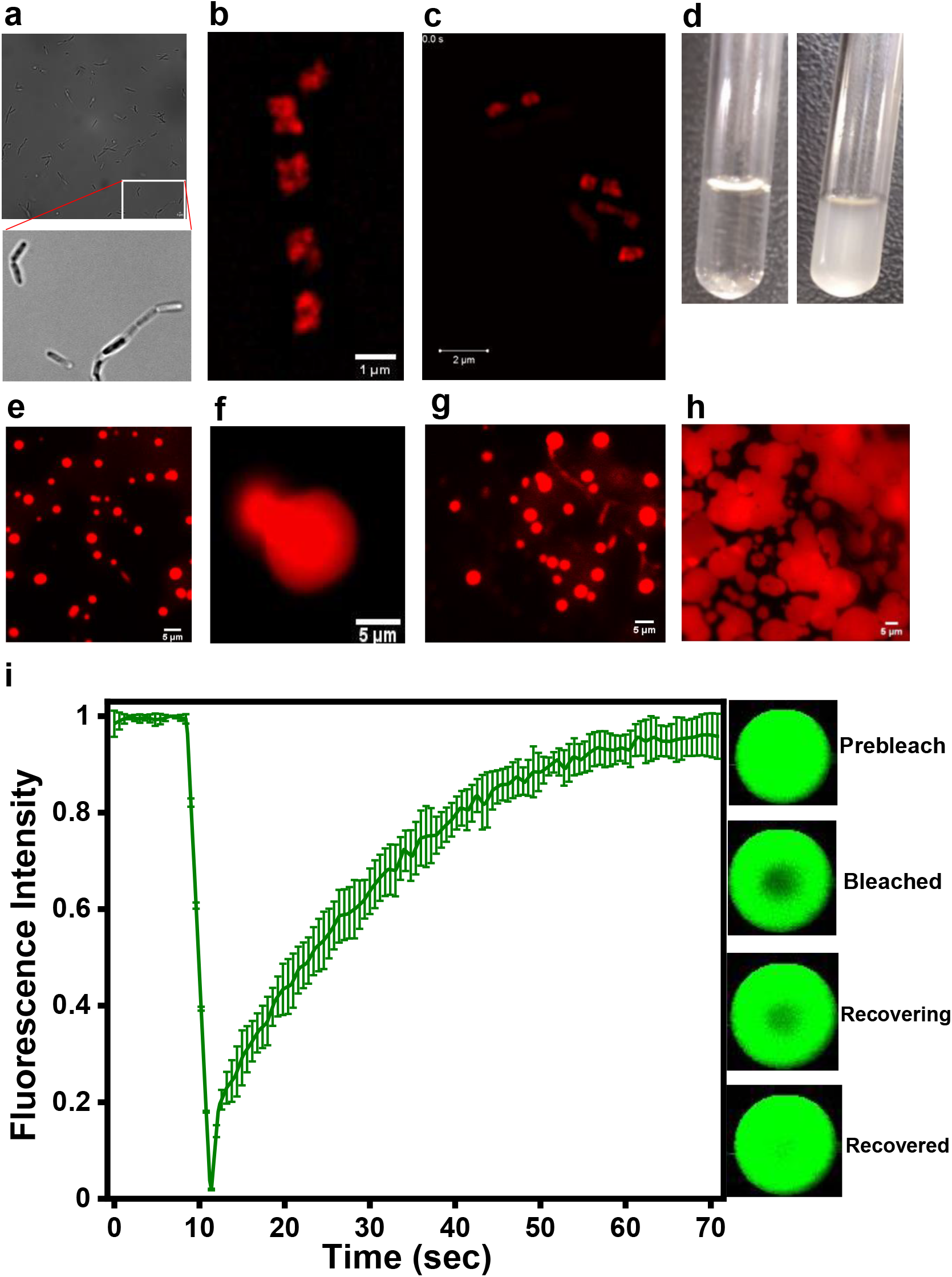
Images of *E. coli* and its nucleoids (displaying dynamic sub-micron-sized domains; panels a-c) and images of phase separation of the NAP, HU-B, into liquid droplet-shaped biomolecular condensates (panels d-m). **a** DIC Image of filaments of unseparated *E. coli* cells. **b** Super-resolution (SIM-derived) image of nucleoids containing sub-micron-sized domains of different shapes in different cells. **c** Single-frame from the SIM videography shown in Supplementary Video 1, demonstrating that the nucleoids’ sub-micron-sized domains are dynamic. **d** Image showing immediate development of turbidity upon addition of 4WJ DNA to HU-B. **e** Confocal fluorescence image demonstrating that the turbidity shown in panel d owes to formation of droplet-shaped condensates of HU-B with 4WJ DNA. **f** Single-frame from a fluorescence video (Supplementary Video 2) demonstrating fusion between droplet-shaped condensates of HU-B. **g** Evidence of dripping from droplets of HU-B. **h** Evidence of surface wetting by droplets of HU-B. **i** Averaged kinetics of fluorescence recovery after photo-bleaching (FRAP) of Alexa 488-labeled HU-B.

### HU causes the accretion of DNA *in vitro*

The likelihood of HU-A, and HU-B, undergoing phase separation was assessed to be significant, based on the distribution of net (linear) charge per residue (NCPR)^41^ in the amino acid sequences of these isoforms, and also based upon a computational prediction of likely regions of disorder, predicted using PONDR^42^ (Supplementary Information Fig. 1). The three-dimensional structures of the single forms of HU that are present in organisms other than *E. coli* are seen to contain regions of structural disorder. Encouraged by HU’s prospects for undergoing phase separation, and with a view to examining whether such phase separation could be triggered by DNA, we examined the effect of adding HU to DNA *in vitro*, by adding unlabeled recombinant forms of HU-A, or HU-B, to salmon testis DNA pre-labeled with a fluorescent DNA-intercalating dye (SYBR Green). The labelled DNA was observed to become instantaneously accreted into compact, spheroidal phase-separated entities upon addition of either HU-A (data not shown), or HU-B (Supplementary Information Fig. 2), over a time scale of seconds (Supplementary Data Video 2).

### HU-B and DNA enter into phase separated liquid droplet condensates *in vitro*

We then examined the formation of phase separated entities containing using HU and a covalently-defined chemical form of DNA already known to bind to HU, instead of salmon testis DNA. We used a synthetic 4-way junction (4WJ) constituted of oligonucleotides associated into a cruciform structure, which we have used previously.^37^ No turbidity was observed in the absence of 4WJ DNA, but instantaneous development of turbidity was observed in HU-B (50 μM) in the presence of 4WJ DNA (3 μM) and a nominal concentration of the macromolecular crowding agent, polyethylene glycol, PEG 6000 (2 %) (Fig. 1**d**). Different modes of imaging established that the turbidity owed to formation of condensates of HU-B and DNA. Bright field, differential interference contrast, phase-contrast, and fluorescence microscopic imaging (Supplementary Information Fig. 4), as well as confocal and/or widefield fluorescence microscopy (Fig.1**e**) established that the droplets formed were spherical. The liquid-like nature of these spherical droplets was then established through observations of droplet-droplet fusion (Fig.1**f**, and Supplementary Video 3), droplet dripping (Fig.1**g**), surface wetting by droplets (Fig.1**h**), and recovery of fluorescence in droplets after photobleaching (FRAP) (Fig.1**i**). Using methods described earlier,^43^ a diffusion coefficient of ~ 2.896 μm^2^ s^-1^ was calculated for HU-B molecules inside these droplets, from the FRAP data (characterized by rapid recovery of ~100 % fluorescence in < 1 min).

Importantly, Supplementary Information Fig. 3 shows that droplets also form when a higher concentration of 4WJ DNA is used (5 μM instead of 3 μM), in the complete absence of any PEG 6000, establishing that HU and DNA together undergo phase separation even when no external agent is used to ‘crowd’ HU macromolecules, through the application of the excluded volume effect. However, although this demonstrated the dispensability of PEG, we chose to use a nominal PEG concentration of 2 % in experiments described hereafter (unless otherwise mentioned) to limit the 4WJ DNA used per experiment to a concentration of 3 μM. Equally importantly, just as HU-B was established to form spherical condensates in the complete absence of PEG when a higher concentration of DNA was used, we also found that HU-B forms spherical condensates in the complete absence of DNA, if a higher concentration of PEG 6000 is used (8 % instead of 2 %), as shown in Supplementary Information Fig. 5. Our view is that these results suggest that the HU-HU interactions responsible for the condensation of HU can involve either (i) a crowding of HU molecules by PEG, or (ii) an accretion of HU molecules that is aided by the accretion of DNA to which HU is bound (i.e., an accretion that is itself triggered by the binding of HU, and the consequent neutralization of charges that lowers repulsions between phosphate groups). The implication of these experiments is that both HU and DNA have the intrinsic ability to undergo phase separation, but that a stimulus is required to reduce their tendencies to remain dissolved in aqueous solution, in order to trigger such phase separation. It is important to note that 4WJ DNA concentrations of both 3 μM and 5 μM are far lower than the 4WJ DNA concentration (60-120 μM) that would adequately represent the base-pair concentration of chromosomal DNA within an *E. coli* cell, corresponding to the physiological presence of ~ 2.25 to ~ 4.50 million base-pairs of genomic DNA in cells of varying size.

### Intermolecular interactions driving HU-B DNA condensation

Having established the liquid-like nature of spherical condensates of HU and 4WJ DNA, we next studied the nature of intermolecular interactions responsible for HU-B’s phase separation. Development of turbidity was used to monitor formation of LLPS droplets per established practice.^7,44^ For temperature-variation experiments, turbidity data alone is shown. For all other variations, microscopy data is additionally shown (Supplementary Information Fig. 6).

#### Effect of varying DNA and protein concentrations

With a (fixed) 50 μM concentration of functional (dimeric) HU-B, turbidity rose progressively with DNA concentration, saturating at ~ 5 μM 4WJ (Fig.2**a**). With a (fixed) 3 μM concentration of 4WJ DNA, turbidity rose progressively with HU-B, saturating above a concentration of ~ 80 μM HU-B (Fig.2**b**). A magnified section of Fig. 2**b** is shown in Supplementary Information Fig. 7, to show that the critical saturation value (C_sat_) of HU-B is ~ 2.5 μM, in the presence of 3 μM 4WJ DNA.

**Figure 2.**
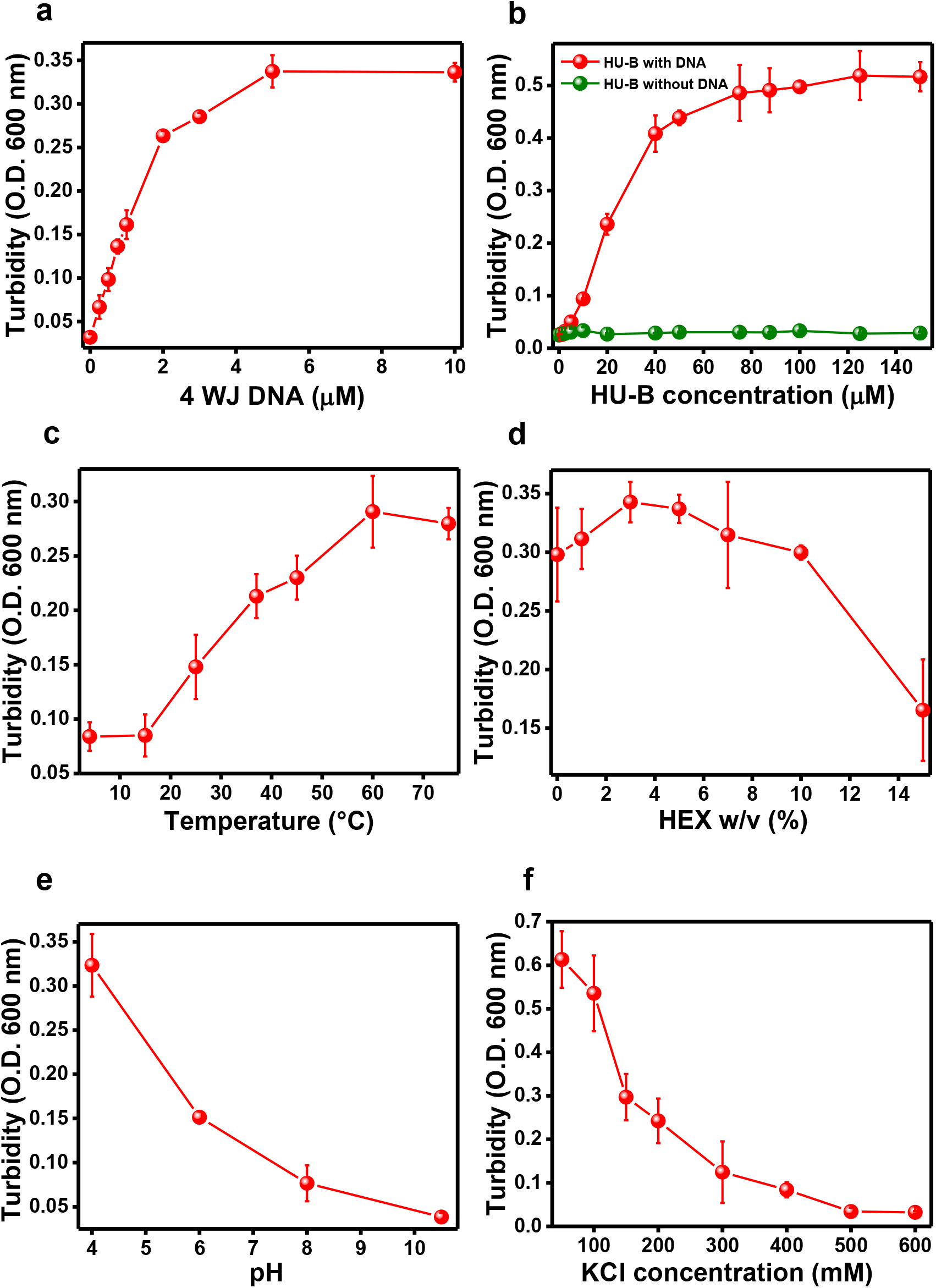
Molecular interactions governing the formation of liquid-liquid phase-separated (LLPS) droplet-shaped condensates of HU-B, assayed through measurement of the immediate development of turbidity, under the following standardized conditions with one parameter varied in each figure panel and others remaining constant: 50 mM Tris; pH 7.4; 150 mM KCl; 50 μM HU-B; 3 μM 4WJ DNA; 2 % macromolecular crowding agent (PEG-6000); 37 ºC. **a** Dependence upon DNA (4WJ) concentration **b** Dependence upon protein concentration (in the presence and absence of DNA). **c** Dependence upon temperature. **d** Response to the presence of HEX. **E** Dependence upon pH. **f** Dependence upon ionic strength (concentration of the salt, KCl).

#### Effect of temperature

Higher temperatures result in higher turbidity (Fig. 2**c**), either due to higher molecular kinetic energies (and the consequent increase in protein-protein collisional frequencies), or through enhancement of hydrophobic contributions to intra-molecular and inter-molecular protein interactions, since hydrophobic interactions would be anticipated to be supported by higher temperatures [especially a specific set of hydrophobic stacking interactions between beta sheets of monomers, in HU-B dimers, or between HU-B and nitrogenous bases of DNA,^45^ exposed through binding of HU-B].

#### Effect of varying the presence of hydrophobic cosolvents

HEX (1,6-hexanediol) disrupts weak hydrophobic interactions relevant to LLPS droplet formation.^46^ A very minor effect was observed, at unusually high concentrations of HEX (Fig. 2**d**), suggesting a minor contribution of weak hydrophobic interactions to the formation of condensates by HU-B and DNA. We have previously shown that hydrophobic interactions are relevant to HU only in respect of interactions between stacked β-sheets at the dimeric interface, which are resistant to disruption by a hydrophobic co-solvent more hydrophobic than HEX, namely dioxane.^45^

#### Effects of electrostatic interactions

There is a rise in turbidity with lowering of pH (Fig. 2**e**), indicating the importance of the number(s) of positive charges upon HU-B in determining its interactions with negatively-charged DNA. As pH is lowered from ~10.0 (HU-B’s pI), there are greater numbers of positive charges upon HU-B, giving rise to greater scope for interaction with DNA, and explaining the increase in turbidity. Interestingly, turbidity also rises with lowering of salt concentration (Fig. 2**f**), presumably because this lowers interactions between HU-B’s positive charges and DNA, through greater levels of ion-exchange mediated by chloride counter-ions (around HU-B’s positive charges) and potassium counter-ions (around DNA’s negative charges).

### HU-A also undergoes LLPS with DNA and forms heterotypic condensates with HU-B

An alignment of the amino acid sequences of *E. coli* HU-A and HU-B (Fig. 3**a**) shows that the isoforms share an identity of ~70 %.^47^ Alignment of structures of monomers of HU-A and HU-B (Fig. 3**b**) yields an RMSD of 0.491 Å. These facts establish that HU-A and HU-B are extremely similar. Therefore, it is reasonable to anticipate that HU-A would also display a tendency to form liquid droplets in the presence of DNA. We discovered that HU-A does indeed form condensates with DNA (at a DNA concentration identical to that standardized for the HU-B experiments, i.e., 3 μM 4WJ). However, HU-A dimer concentrations are required to exceed ~45 μM for such condensation to occur (Fig. 3**c**), revealing that HU-A dimers are eighteen times poorer at forming condensates with 3 μM 4WJ DNA than HU-B dimers, earlier established to have a C_sat_ concentration of ~2.5 μM (Supplementary Information Fig. 7). Unsurprisingly, condensates of HU-A with DNA also proved to be liquid-like, displaying rapid recovery of fluorescence in FRAP experiments (Fig.3**d**). Our data thus suggests that both HU-A and HU-B form spherical condensates with DNA; only at different protein concentrations. Phase diagrams presenting effects upon HU-A/HU-B turbidity of alterations in salt concentration (Fig. 3**e**), or protein concentration (Fig. 3**f**), further establish that HU-A is indeed much poorer than HU-B at forming condensates with DNA. Importantly, we found that HU-B and HU-A form heterotypic condensates^48^ with each other, and with DNA, when all three species are present together (Fig.3**g**). This suggests that the ratio of HU-A:HU-B in a particular region of the bacterial chromosome could determine the extent to which that region exists in a state of phase separation, since the abilities of HU-A and HU-B in regard of causing phase separation differ by a little over one order of magnitude.

**Figure 3.**
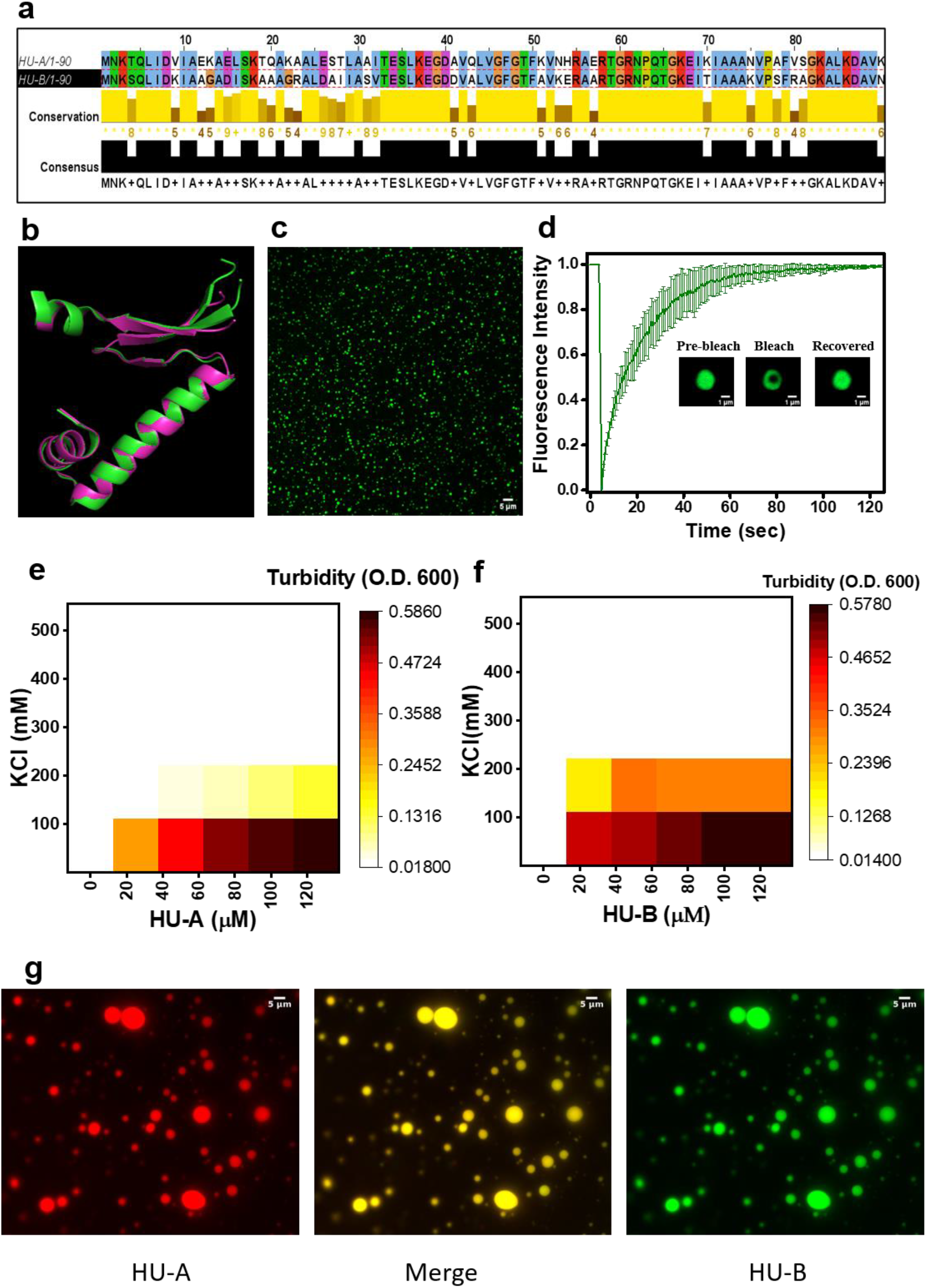
Comparison of the sequences, structures, and phase separation behavior of HU-A and HU-B, and demonstration of their ability to coacervate with each other. **a** Alignment of the amino acid sequences of HU-A and HU-B. **b** Alignment of the structures of the polypeptide chains of HU-A (green; PDB ID 1MUL) and HU-B (magenta; PDB ID 4P3V). **c** A representative image showing small droplet-shaped condensates of HU-A (110 μM). **d** Fluorescence recovery after photo-bleaching (kinetics and images) of Alexa 488-labeled HU-A in droplet-shaped condensates. **e** Phase diagram of HU-A as function(s) of salt and protein concentrations. **f** Phase diagram of HU-B as function(s) of salt and protein concentrations. **g** Complex coacervation or of HU-A and HU-B into the same droplet-shaped condensates.

### The degree of condensate formation appears to be correlated with HU’s multimericity

It has been speculated that a higher degree of multimericity translates into a greater valency in respect of intermolecular interactions, with this further (positively) affecting the formation of LLPS condensates.^49^ With the isoforms of HU, it is reported that HU-A exists mainly in the form of dimers, whereas HU-B exists in the form of dimers, tetramers, and octamers.^50^ We re-established this behavior, using the recombinant proteins deployed in the current experiments, in three different ways: (i) In gel filtration (size exclusion) chromatography experiments, HU-A was observed to elute as a single (dimeric) peak, whereas HU-B eluted as a mixture of species of sizes ranging from dimer to octamer (Fig.4**a**); (ii) In acidic native polyacrylamide gel electrophoresis (PAGE) experiments, HU-A migrated as a single electrophoretic band, whereas HU-B migrated as a smeared band, consistent its presumed interconversion between multimeric states during electrophoresis (Fig.4**b**); (iii) In, electrophoretic mobility shift assays (EMSA) examining the binding of HU-A, or HU-B, to 4WJ DNA, HU-A generated discrete mobility-shifted species for HU-A-DNA complexes (Fig.4**c**), whereas HU-B generated both such species and also additional DNA-bound states incapable of entering the gel (Fig.4**d**), consistent with HU-B’s formation of larger complexes with DNA, due to its higher multimericity. To examine whether these differences in multimericity are correlated with the different C_sat_ values of HU-A (~45 μM) and HU-B (~2.5 μM), we generated an HU-A mutant incorporating two point mutations, E34K and V42L. This mutant is reported to form dimers and also octamers, but no tetramers,^50^ unlike HU-A (entirely dimeric), and HU-B (dimeric, tetramic, and also octameric). Turbidity plots for HU-A, HU-B, and E34K:V42L HU-A, show that there is a dramatic down-shifting of C_sat_ value from HU-A (~45 μM), to E34K:V42L HU-A (>10 μM), to HU-B (~2.5 μM). Confocal microscopic imaging also confirmed that the E34K:V42L HU-A’s propensity to form spherical condensates is intermediate to those of HU-A and HU-B (Fig.4**f**).

**Figure 4.**
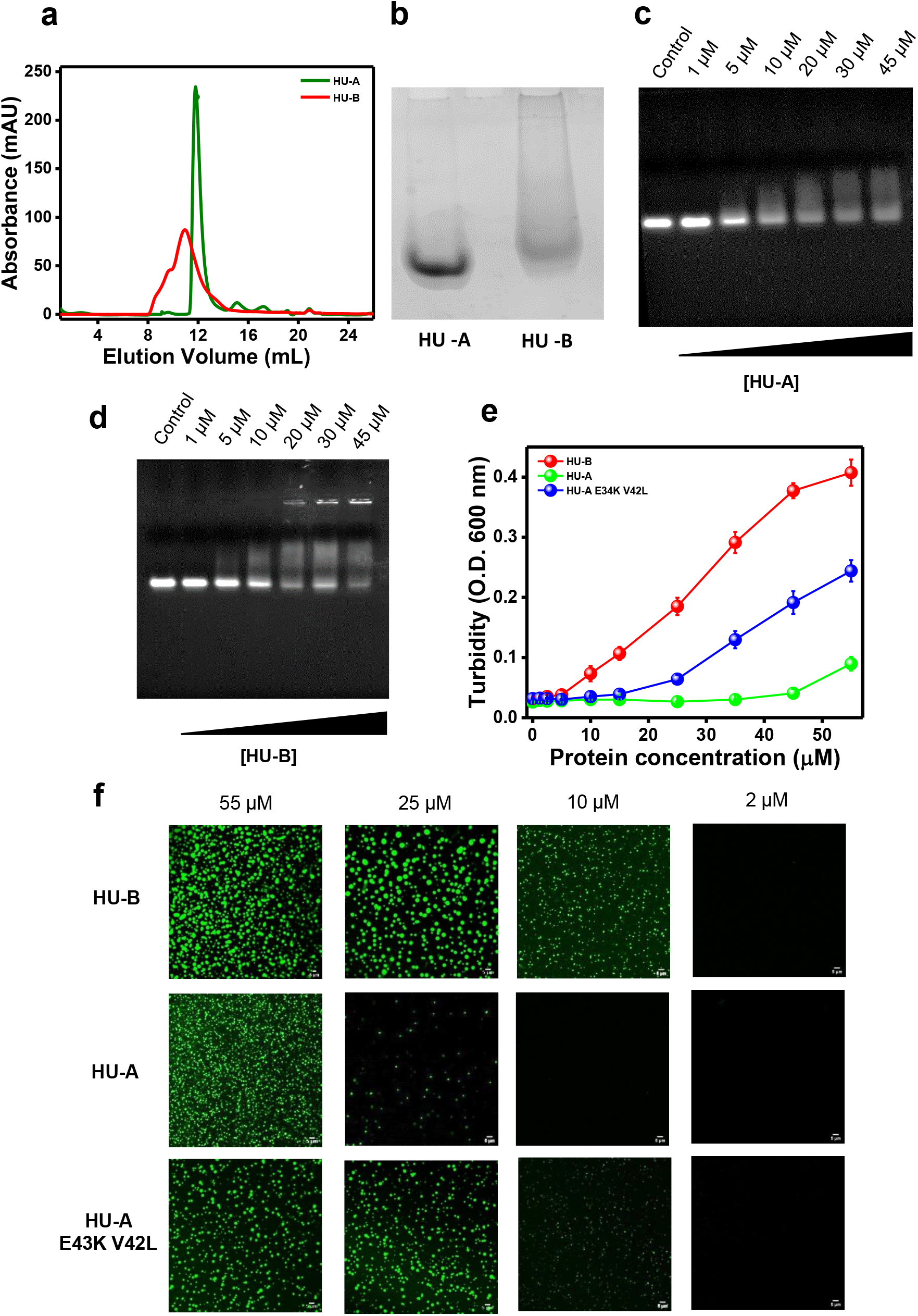
The role of multimericity in the phase separation behavior of HU. **a** Size exclusion chromatogram of HU-A (green) and HU-B (red). **b** Electrophoretic behavior of HU-A and HU-B on native PAGE gels (lanes as marked). **c** EMSA assay of HU-A binding to 4WJ DNA (1 μM). **d** EMSA assay of HU-B binding to 4WJ DNA (1 μM). **e** Turbidometry plots showing differences in the c_sat_ values of HU-A, E34K-V42L HU-A, and HU-B. **f** Panels showing confocal microscopic images of condensate number(s) and size(s) of HU-A, E34K-V42L HU-A, and HU-B, at different protein concentrations.

It is known that a heterodimer of HU-A and HU-B dominates the log phase of growth of *E. coli* cultures.^51^ We examined the condensate-forming behavior of HU-B/HU-A heterodimers, by examining the behavior of a proxy, or simulacrum, of such a dimer, created through a genetic fusion of HU-B and HU-A, as described by us earlier.^52^ The concentration-dependent turbidity plot of this HU-B/HU-A simulacrum (Supplementary Information Fig. 8**a**) shows that it also forms condensates with DNA under conditions and concentrations similar to those used to examine condensate formation by HU-B (Supplementary Information Fig. 8**b**). This establishes that HU in all of its homo-dimeric and hetero-dimeric forms engages in the formation of liquid droplet condensates with DNA.

### Dps also undergoes LLPS and forms multiphasic condensates with HU-B and DNA

Dps, which is acidic, and dodecameric, is a major NAP of the stationary phase of *E. coli* and other bacteria. It interacts with DNA through a basic N-terminal tail present in each monomer. It also contains some intrinsically disordered regions (IDRs) and charge clusters throughout its sequence (Supplementary Fig. 9**a**,**b**).^53,54^ Fascinatingly, the already well-known enhancement of compaction of the nucleoid during the stationary phase of growth of *E. coli*, happens to be correlated with the increased expression of Dps, and the reduced expression of HU-A and HU-B.^23,55^ Stress is known to induce phase separation of Dps into crystalline and liquid-like crystalline structures *in vivo*.^54,56–58^ Dps is also known to phase-separate into crystals *in vitro*.^54,56^ The concentration of Dps monomers/cell rises to 1,80,000 (~15,000 dodecamers),^55^ translating into a bacterial cell volume-dependent concentration of 18.75-50.0 μM. We tested the ability of 30 μM Dps to form spherical condensates, in the absence, and also in the presence, of DNA, HU-A and/or HU-B. Dps showed phase separation with Salmon testis DNA (Fig.5**a**), forming condensates smaller than those formed by HU, with evidence of formation of both spherical and non-spherical entities (appearing to result, in some instances, from irregular adherence of spherical condensates without undergoing complete fusion). Dps condensates displayed rapid FRAP (Fig.5**b**). Despite the non-canonical (i.e., non-spherical) appearance of some of these condensates, confocal microscopic imaging confirmed the co-localization of Dps protein and Salmon testis DNA (Fig. 5**c**).

**Figure 5.**
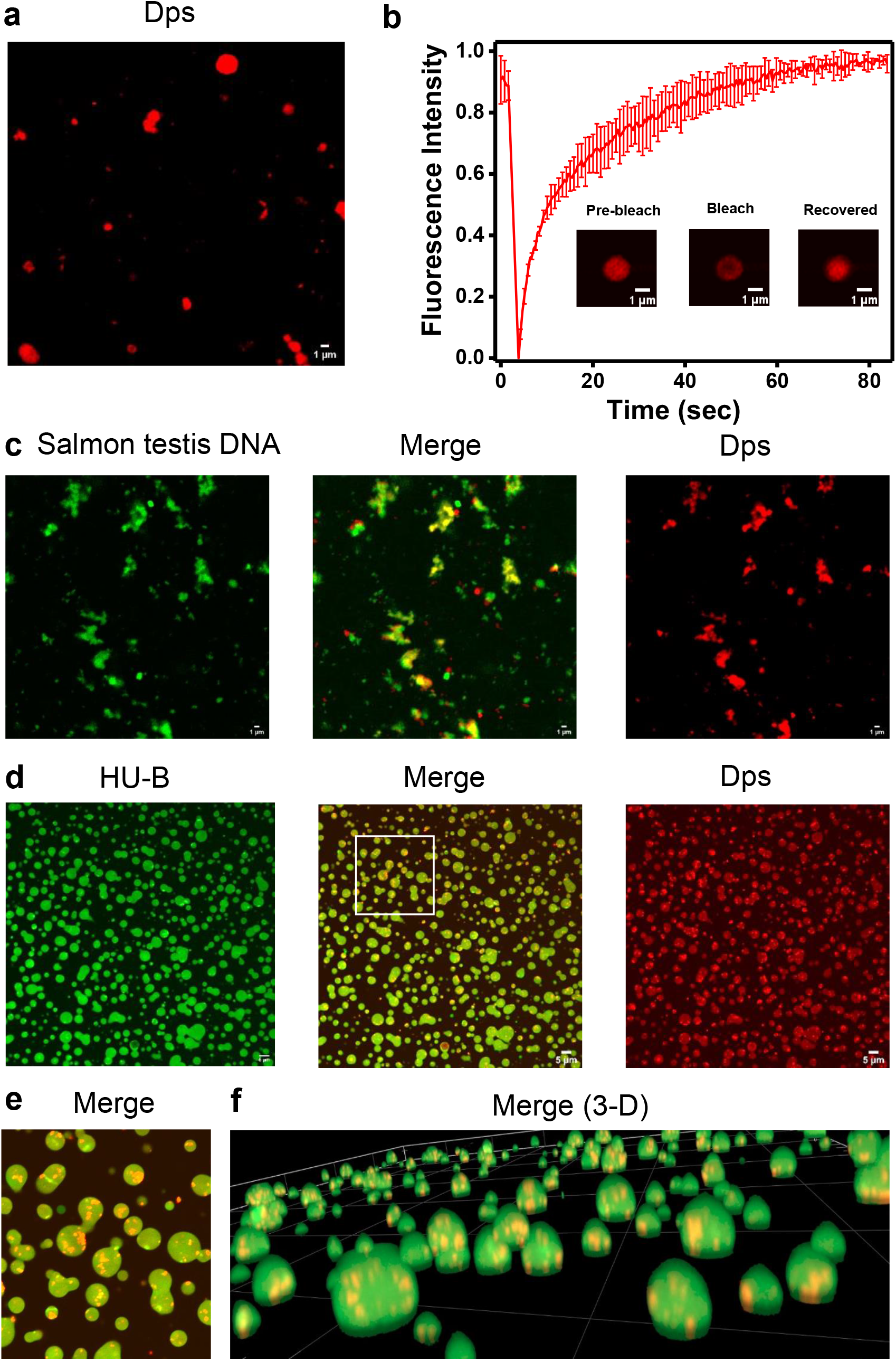
Phase separation behaviour of Dps with DNA, in the absence and presence of HU-B. **a** Representative confocal microscopic image of condensates of Dps. **b** Kinetics and images of fluorescence recovery after photo-bleaching (FRAP) of Alexa 594-labeled Dps. **c** Salmon testis DNA labelled by SYBR Green (green), Dps labelled by Alexa 594 (red), and merged channels, for condensates of Salmon testis DNA and Dps. **d** HU-B labelled by Alexa 488 (Green), Dps labelled by Alexa 594 (red), and merged channels, for condensates of HU-B, unlabelled 4WJ DNA and Dps. **e** Enlarged section of image shown in panel d showing Dps-rich droplets containing more Dps than HU-B, enclosed within HU-B-rich droplets containing more HU-B than Dps. **f** A three-dimensional confocal image of the HU-B, 4 WJ DNA and DPS droplets similar to panel d.

Additionally, we confirmed that Dps forms LLPS condensates by itself, in the absence of DNA, when 8 % PEG 6000 is present (Supplementary Information Fig. 9**c**). Upon addition of DNA to solutions containing Dps and HU, Dps co-localized within condensates of HU-B (Fig. 5**d**), as well as HU-A (Supplementary Figs. 10), becoming incorporated within such condensates in the form of irregularly-shaped Dps-rich domains of smaller volume. This result may be viewed in light of the speculation that some proteins can act as client proteins in scaffolds prepared by other proteins, during phase separation.^59^ What is remarkable about these condensates of HU-B (green), 4WJ DNA, and Dps (red), is the evident clustering of Dps-rich droplets (orange) within larger HU-rich droplets (green), indicative of the formation of phases of different grades of liquidity that do not manage to mix fully (Fig. 5**d**). A magnified image of one such group of droplets-within-droplets (Fig. 5**e**) and a three-dimensional cross-section of a field containing many droplets-within-droplets (Fig. 5**f**) are also shown. Interestingly, when HU-B and Dps were subjected to phase separation in the absence of DNA, through inclusion of 8 % PEG, uniform mixing was observed, giving rise to spherical condensates containing no regions that were distinguishably rich in either HU-B, or Dps (Supplementary Information Fig. 11). This suggests that the irregular shapes of Dps-DNA condensates, or of Dps-dominated (HU-B-Dps-DNA) condensates enclosed within spherical HU-B-dominated condensates, could be related to the manner in which Dps binds to DNA, and organizes it. The summary view would be that HU-B, and Dps, two major NAPs present in stationary-phase *E. coli*, are capable of co-existing within the same condensates. The possibility of HU-B and DNA forming a different phase within nucleoid interiors, from the phases formed by mixtures of Dps, DNA, and HU-B, or HU-A, at nucleoid surfaces, has also not escaped our notice. The three protein species seem to offer a near-continuum of extents of phase separation, and compatibility with cytoplasm.

### HU-B undergoes LLPS with multiple forms of nucleic acids and DNA polymerase

We explored the ability of HU-B to undergo condensation/coacervation with different forms of DNA/RNA, using experiments simulating the situation inside a nucleoid where different forms of DNA and RNA, and different proteins like HU-B, HU-A and Dps, exist together with other NAPs, and DNA-binding proteins, e.g., DNA polymerase. Fig. 6 shows independent and merged images of differentially-labeled forms of HU-B with different combinations of HU-A, Dps, 4WJ DNA, nicked DNA, double-stranded DNA, and single-stranded DNA, showing the co-localization of multiple and varied species into the same droplets. We also found HU-B to phase separate with Poly-U RNA (Supplementary Information Fig. 12). Notably, knockout of a non-coding HU-binding RNA lead to de-compaction of the nucleoid.^60^ Lastly, we examined whether DNA polymerase can exist within condensates of HU and DNA, through an experiment in which fluorescently-labeled Pfu DNA polymerase was mixed with condensates of DNA, and Venus-HU-B.^37^ DNA polymerase was observed to co-localized with HU-DNA condensates (Supplementary Information Fig. 13), suggesting that HU-mediated phase separation of genomic DNA would be compatible with the presence of other DNA-binding protein species.

**Figure 6.**
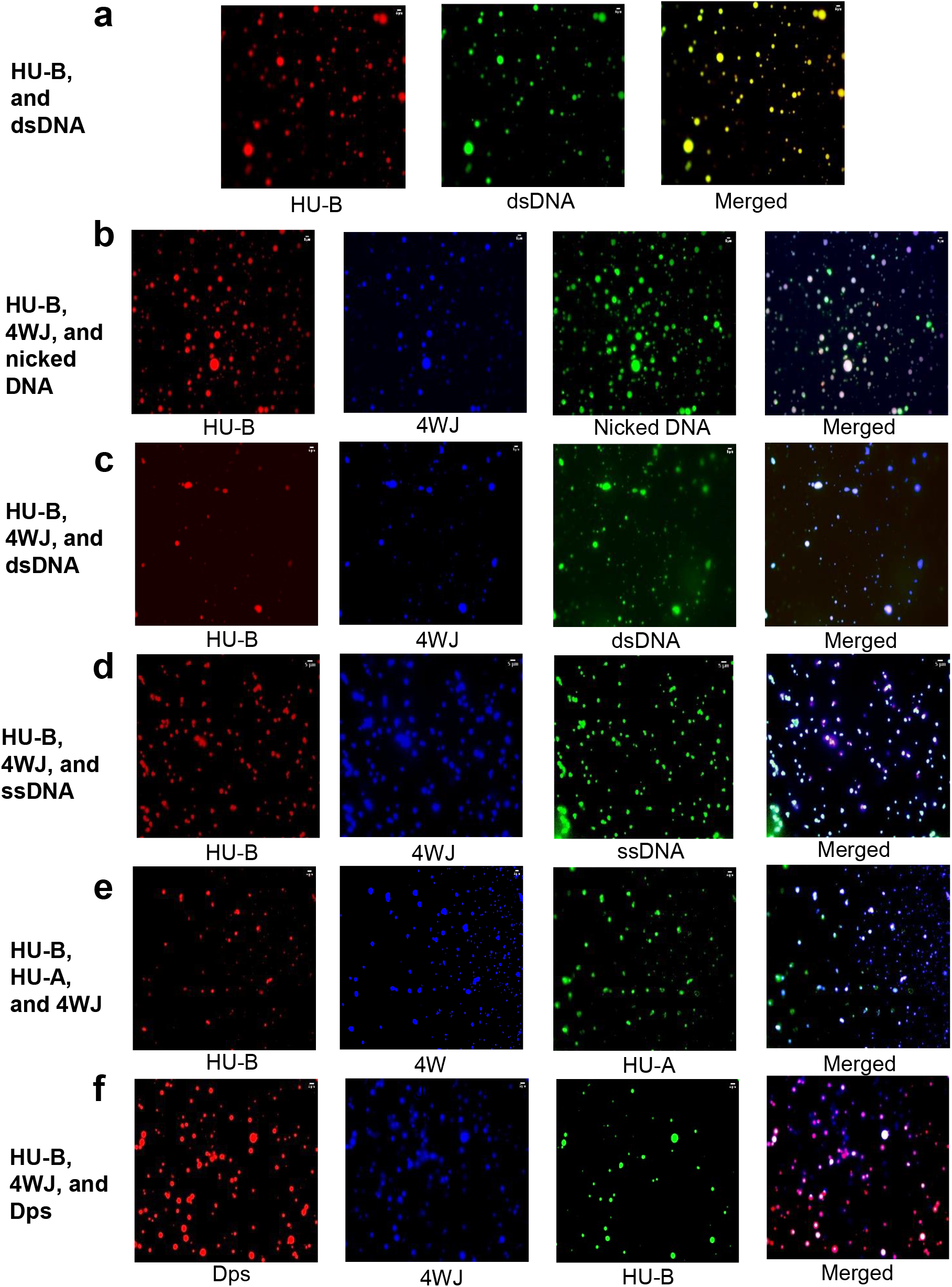
Coacervation or co-phase separation of HU-B (labeled by Alexa 594, or Alexa 488) with multiple different forms of DNA (labeled by DAPI, SYBR Green, or 6-fluorescein amidite), and with other NAPs, such as Dps (labeled by Alexa 594) or HU-A (labeled by Alexa 488). **a** Coacervates of HU-B (red) and ds DNA (green). **b** Coacervates of HU-B (red), 4WJ DNA (blue), and nicked DNA (green). **c** Coacervates of HU-B (red), 4WJ DNA (blue), and ds DNA (green). **d** Coacervates of HU-B (red), 4WJ DNA (blue), and ss DNA (green). **e** Coacervates of HU-B (red), 4WJ DNA (blue), and HU-A (green). **f** Coacervates of Dps (red), 4WJ DNA (blue), and HU-B (green).

## Discussion

The idea that the nucleoid is phase-separated from the cytoplasm in bacteria has existed for decades.^22^ In recent years, evidence has also accumulated for liquid-liquid phase separation of various proteins in the company of nucleic acids, e.g., HP1 in heterochromatin,^61^ NPM1 in the nucleolus,^62^ H1 in nucleosomes,^63^ and TFAM in mitochondrial nucleoids.^17^ In bacterial nucleoids, HU and Dps, which are the most abundant of *E. coli*’s NAPs, have been proposed to be involved in the formation of a phase-separated state;^64^ however, without any experimental evidence.

*E. coli*’s genomic DNA (2 mm in length) is compacted into a tiny volume (~ 0.5-3.0 femtolitres). Within this volume, DNA remains engaged in replication and transcription, within cells of tiny dimensions (~ 0.5 – 3.0 μm). NAPs must play a critical role in the accretion and compaction of DNA into such tiny volumes, while helping to overcome the mutually-repulsive interactions of phosphate groups, through neutralization of DNA’s charges. Coacervation of proteins and DNA into phase-separated states potentially allows DNA-protein interactions to assist in the process of DNA compaction, with abundant and non-specifically DNA-binding proteins managing to do this over entire chromosomes, driving a form of self-crowding behavior through charge neutralization.

Since HU has a basic pI, it could potentially organize DNA into LLPS states in nucleoid interiors, where HU could remain buried by bound DNA. Since Dps has an acidic pI (and binds to DNA through a basic N-terminal tail), it could potentially organize DNA into LLPS states by itself largely at the nucleoid-cytoplasmic interface, as charge-charge repulsions with DNA would prevent Dps from becoming completely buried. However, HU could also engage in the neutralization of the negative overall charge on Dps in a non-specific manner. Thus, it is our view that HU and Dps constitute a ‘minimalistic’ set of proteins for phase separation of the bacterial chromosome into a nucleoid.

Our findings reveal that HU-A and HU-B, form spherical heterotypic condensates with double-stranded DNA as well as with other forms of nucleic acids, and we show that HU’s interaction with DNA causes DNA to be accreted under physiological conditions of temperature, pH, and ionic strength, and at physiological HU concentrations, with sub-physiological concentrations of nucleic acids, in the absence of other macromolecular crowding agents. Thus, even higher (physiological) DNA concentrations and the presence of other macromolecules could cause HU and DNA to engage in LLPS at even lower local concentrations of HU than demonstrated by us (~2.5 μM HU-B, or ~50 μM HU-A). We also show that Dps engages in LLPS with DNA, and that HU and Dps together form complex multiphasic droplets with DNA, without undergoing complete mixing, and with evidence of formation of domains of differential HU or Dps content. This ability of HU condensates to accommodate Dps-rich condensates could prevent the premature crystallization of Dps during stress. In summary, we show that HU A/B and Dps influence each other, and the nucleoid, through phase separation, in a manner that could be ‘tuned’ by stress for growth phase regulation in *E. coli* cultures (Fig. 7). Our super-resolution imaging results also suggest that nucleoids are dynamic assemblages of HU-containing liquid droplets. Notably, it has been reported that genetic double-knockouts of HU-A and HU-B leads to de-compaction of the nucleoid.^65–67^

**Figure 7.**
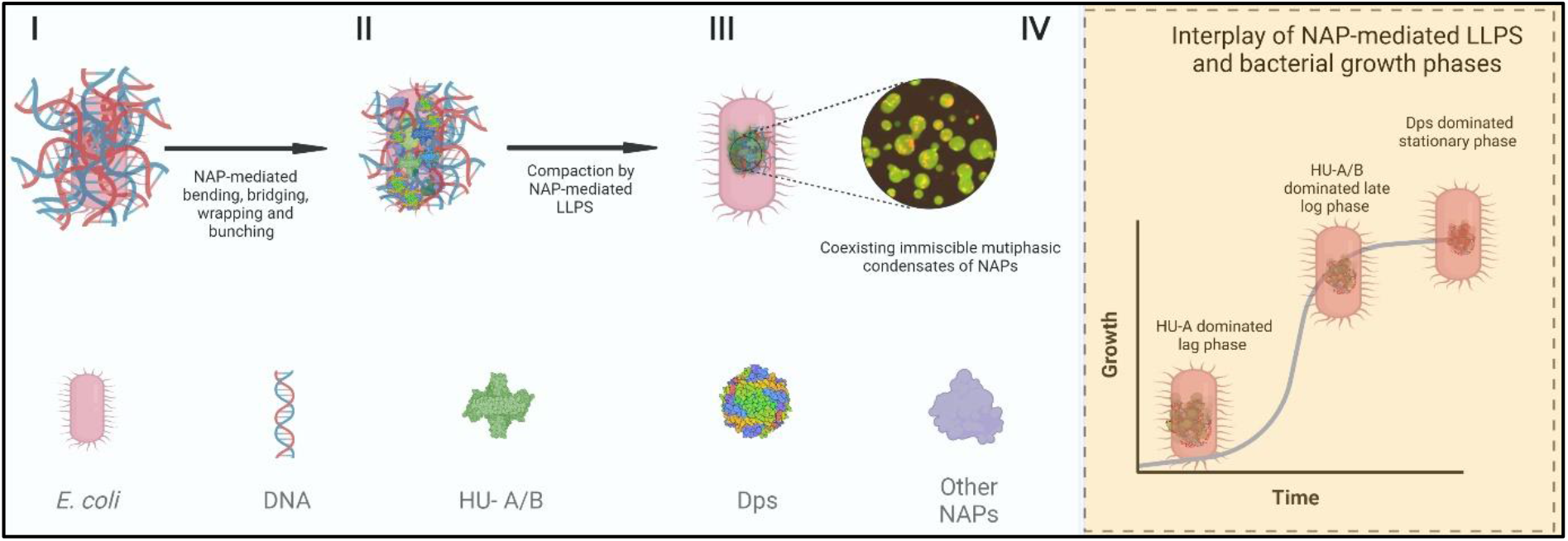
Model showing role of NAPs and LLPS in the compaction of the *E. coli* genomic DNA. I) Uncompacted DNA cannot be contained within *E. coli*. II) DNA is significantly compacted by NAP-mediated bending, bridging, wrapping and bunching, but not sufficiently to allow the over two thousand-fold compaction required. III) NAP-mediated LLPS assists in further compaction and in the maintenance of a state of compaction, with sub-micron-sized droplets fusing and separating as required, while adopting different grades of liquidity through differential presences of different NAPs. The expansion of a section of the nucleoid shows multiphasic condensates of NAPs, (taken from Fig 5**e**). IV) Low level of HU-A homodimer-mediated LLPS in the lag phase of growth; Higher level of HU-B homodimer-mediated and HU-A/B heterodimer-mediated LLPS in the late log phase of growth; Highest level of HU-B homodimer-mediated and Dps-mediated LLPS in the stationary phase, along with liquid-solid phase separation of Dps into a crystalline or semi-crystalline state.

Taken together, our work suggests that the nucleoid comprises sub-micron-sized, spherical phase-separated multiphasic condensates, connected by a thread of DNA that runs through such droplets like a ‘bundled-up necklace’, resolvable into independent phase-separated beads and also fusible into larger beads, based on the state of DNA supercoiling, bending, looping, and binding to HU-B, HU-A, Dps, and other NAPs. Recently, single particle-tracking of a photoactivable form of HU has shown that it displays two types of behaviors within nucleoids; one showing high mobility, and the other showing restricted mobility.^68^ We propose that this is consistent with different coexisting immiscible liquid phases of differing density within *E. coli* nucleoid, with some HU populations packed away with DNA remaining largely buried between cell generations, and other HU populations associated with the cytoplasm and regions undergoing replication, or transcription. These findings highlight the first evidence of phase separation in bacterial nucleoids and can serve as a model for the study of bacterial regulation through the lens of NAP-mediated phase separation of bacterial nucleoids. Similar immiscible multiphasic nucleoplasmic condensates may be found across all organisms in future studies, and serve as a stepping-stone for the synthesis of artificial organelles for synthetic biology applications.

## Methods

### Nucleic acids

#### Synthetic oligonucleotides

All oligonucleotides (oligos) were purchased from Sigma-Merck and were resuspended in Tris buffer at pH 7.5.

#### Fragmented genomic DNA

Salmon testis DNA was purchased from Sigma-Merck (Cat no. D-9156).

#### Poly(U) RNA

Poly(U) RNA purchased from Sigma-Merck (Cat no. P9428).

#### Single-stranded DNA

The oligo, 5’-TGATCATGCATCGTTCCACTGTGTCCGCGACATCTACGTC-FAM - 3’ was used as single-stranded (ss) DNA.

#### Double-stranded DNA

Double-stranded (ds) of a length of 20 basepairs was prepared by annealing two oligos, 5’-GTTCAATTGTTGTTAACTTG-3’ and 5’-CAAGTTAACAACAATTGAAC-3’.

#### Nicked double-stranded DNA

Nicked ds DNA was created by annealing the long oligo 5’-GACGTAGATGTCGCGGACACAGTGGAACGATGCATGATCAGCAAGGACGATCCTGTCTTGGTGGTAAGGGTGCGC-3’ to two short oligos, 5’-TGATCATGCATCGTTCCACTGTGTCCGCGACATCTACGTC-FAM-3’ and 5’-GCGCACCCTTACCACCAAGACAGGATCGTCCTTGC-3’.

#### Four-way junction (4WJ) DNA

4WJ DNA was generated through the annealing of 4 different oligos in equimolar amounts to create a Holliday junction-like (cruciform) structure. The four oligos used were:

5’-CCCTATAACCCCTGCATTGAATTCCAGTCTGATAA 3’,

5’-GTAGTCGTGATAGGTGCAGGGGTTATAGGG-3’,

5’-AACAGTAGCTCTTATTCGAGCTCGCGCCCTATCACGACTA-3’,

5’-TTTATCAGACTGGAATTCAAGCGCGAGCTCGAATAAGAGCTACTGT-3’.

#### DNA fluorescent labeling

4WJ DNA and ds-DNA were fluorescently-labeled with DAPI and SYBR green respectively, free fluorophores were removed through repeated ultrafiltration and washing.

### Construction of genes encoding wild-type or mutant HU/Dps

Strategies for the cloning of fusion constructs involving the fluorescent proteins RFP, or Venus, placed at the N-termini of HU isoforms (with the intent of creating either RFP-HU A, or Venus HU-B) have already been described.^37^ In the same descriptions of work, strategies for the cloning of an F47W mutant of HU-B, and an F79W mutant of HU-A have also been described, together with the creation of a fusion of HU-B and HU-A (HU Simulacrum) incorporating an 11 amino acids-long flexible GS linker placed between the two proteins. We replaced the wild-type HU-B and HU-A in the HU simulacrum with the said F47W mutant of HU-B, and the F79W mutant of HU-A, respectively, to create F47W-HU-B and F79W-HU-A, using standard recombinant DNA techniques. In further developments of mutants, we used the genetic background of the F47W mutation in HU-B, or the F79W mutation in HU-A, to carry out the requisite site-directed mutagenesis, using splicing by overlap extension (SOE) PCR-based strategies entirely identical to those described earlier,^37^ as well as mutation-specific primers designed to create HU-B or HU-A genes encoding the following mutations: (i) an N-terminal cysteine in HU-B, (ii) an S35C mutation in HU-B, (iii) an S17C mutation in HU-A, (iv) an E34K-V42L double mutation in HU-A already containing the F79W mutation. Separately, an S22C mutation was created in Dps which already contains 2 tryptophan residues and does not require the introduction of a tryptophan (W) residue as a spectroscopic reporter (unlike HU-B or HU-A). The gene encoding Dps was cloned by amplification from the *E. coli* genome through a PCR reaction in which the forward primer contained an Nde I restriction site, and the reverse primer contained an Xho I restriction site, with the genomic DNA template sourced through lysing of whole *E. coli* cells by heat (98 °C, 5 min) immediately prior to 35 cycles of PCR [Denaturation: 95 °C, 30 sec); Annealing: 55 °C, 45 sec; Extension: 72 °C, 60 sec] using the Go-Taq Flexi (Promega) thermostable DNA polymerase. The amplicon was extracted from the agarose gel, digested and ligated between Nde I and Xho I sites of the pET-23a vector, by T4 DNA ligase. DNA sequences of all clones of HU and Dps were confirmed through sequencing by dideoxy chain termination, using a commercial service (Agrigenome).

### Protein expression and purification

#### HU-A and HU-B

Genes encoding HU-A and HU-B, and mutants bearing 6xHis affinity tags at their N-termini, were cloned into the pQE-30 vector (between the *Bam HI* and *Hind III* restriction sites) for expression in either the XL-1-Blue strain or the M15 (pRep4-carrying) strain of *E. coli*. HU-B and its mutants were expressed in the XL1-Blue strain, whereas variants of HU-A were expressed in the M15 strain. Cultures were grown for 8 h, and cells were harvested through pelleting by centrifugation at 9000 *g*. Pelleted cells were re-suspended in phosphate-buffered saline (PBS) supplemented with 1 M NaCl, to facilitate separation of HU from DNA (and, consequently, also DNA-bound proteins) during subsequent purification. The re-suspended cells were lysed using a sonicator (Qsonica/Misonix). Removal of cell debris was done through centrifugation at 15000 *g* for 1 h. Supernatants were subjected to Ni-NTA immobilized metal affinity chromatography (IMAC), using standard conditions (Qiagen). Protein purified through Ni-NTA chromatography was subjected to further (polishing-step) purification, using cation exchange chromatography performed on either a HiFliQ S-type (Protein Ark) column or a Bio-Rad (Econo) column packed with Mono-S resin (GE). The purified protein was then dialyzed against 50 mM Tris buffer of pH 7.4, containing 150 mM KCl, concentrated to ~3 mg/ml, partitioned into aliquots, flash-frozen in liquid nitrogen, and stored at −80°C. *<u>Dps</u>*. Dps and its cysteine-containing variants were cloned into the pET-23a vector between the *Nde I* and *Xho I* restriction sites, for expression in the BL21 (DE3) pLysS* strain of *E. coli*, through induction by IPTG (1 mM) at a culture O.D of 1.0, followed by 24 h of growth of culture(s). The method for purification of C-terminally 6xHis-tagged Dps through Ni-NTA (IMAC) chromatography was identical to that used for HU. Since high purity was obtained using IMAC, Dps and its variants weren’t subjected to ion-exchange chromatography, as was required with HU-B or HU-A.

### Estimation of HU/Dps concentration

Protein concentrations were estimated using the 280 nm absorption of tryptophan, and estimated extinction coefficients. Native HU-A and HU-B do not contain the residue tryptophan. Since a concentration-dependent phenomenon like phase separation cannot be studied with precision using tryptophan-lacking proteins (since the estimation of concentration proves to be difficult), we first verified, using roughly-estimated concentrations, that wild-type HU-B and HU-A display LLPS behavior. Thereafter, we performed standard experiments only with the F47W mutant of HU-B, or the F79W mutant of HU-A, with an accurate estimation of protein concentrations based on the tryptophan residue present in these mutants, which have previously been used and shown to be identical to wild type HU protein in respect of DNA binding.^37^

### Electrophoretic mobility shift assay (EMSA)

Protein samples of varying concentrations were mixed with 4-way junction (4WJ) DNA to yield a final DNA concentration of 1 μM, and loaded onto a 0.5 % agarose gel in TAE buffer, using a constant voltage of 60 V, for 1 h. Protein-bound DNA and free DNA on the gel were then stained with ethidium bromide (EtBr) and visualized through UV-induced fluorescence.

### Phase separation assays

First standard physical/chemical conditions, and reagent concentrations, were established, as described in the next six subsections. In the seventh subsection, a description is provided of the ranges over which each of these physical/chemical conditions, or reagent concentrations, was varied, while maintaining all the others at a fixed value.

#### (1) pH and ionic strength

For examining the ability of coacervates of protein and DNA to form LLPS droplets, a standard pH of 7.4 (50 mM Tris buffer), and a standard salt concentration of 150 mM KCl, were chosen, with the intent of mimicking the pH and ionic strength of the bacterial cytosol. A temperature of 37 °C was chosen, with the intent of mimicking the temperature at which a culture of a bacterium such as *E. coli* grows optimally.

#### (2) HU concentration

A standard HU dimer concentration of 50 μM (i.e., HU monomer concentration of 100 μM) was chosen, with the intent of mimicking an HU concentration of ~1 mg/ml for LLPS experiments, corresponding to 60,000 HU monomers per femtolitre, since the average physiological concentration of HU varies from 30,000 to 60,000 monomers per cell, and since cells display variation of volume from 0.5 to 3.0 femtolitres.

#### (3) Dps concentration

A standard Dps dodecamer concentration of 30 μM (i.e., Dps monomer concentration of 360 μM) was chosen to mimic the cellular concentration of Dps (12.5 μM for a cell of 2 femtolitre volume, and varying between 9.38 μM and 50 μM for cell volume with volumes varying from 0.5 to 3.0 femtolitres; assuming 180,000 Dps monomers per cell).

#### (4) 4WJ DNA concentration

A cruciform-shaped DNA Holliday 4-way junction (4WJ; made up of ~110 base pairs) concentration of 3 μM (~0.15 mg/ml) was chosen to create a DNA concentration of 112,500 base pairs per femtolitre. This DNA concentration happens to under-represent, by a factor of 20 to 40, the average physiological concentration of ~ 2.25 to ~ 4.50 million base-pairs per femtolitre (corresponding to one half-genome, or a full genome per femtolitre).

#### (5) ssDNA and plasmid DNA concentrations

Single-stranded salmon testis DNA with an average length of ~ 1200-1600 bases, and linearized plasmid DNA with a length of 4.2 kilobase-pairs, were used at a concentration of 0.15 mg/ml.

#### (6) Crowding agent concentration

After confirming that LLPS behavior is also seen in the absence of a crowding agent, a standard (nominal) PEG-6000 concentration of 2 % (w/v) was chosen to under-represent concentrations of this crowding agent (5-20 %) which are commonly used to simulate cellular conditions and the crowding of macromolecules in the cell cytoplasm.

#### (7) Variations of conditions and concentrations

These were employed to map the ranges of conditions under which HU-B forms LLPS droplets, by varying one condition at a time (e.g., temperature, pH or ionic strength) while maintaining all other standardized conditions (described above). A single readout of turbidity was measured after confirming fully that the turbidity reflects the formation of LLPS droplets. To examine the effect of varying chemical conditions, e.g., pH or ionic strength, turbidity measurements were made immediately after mixing of protein with DNA, using a wavelength of 600 nm, on a 96-well plate (Eppendorf cat no. 0030730011) in a plate reader (BMG Labtech POLARstar Omega). For examination of the effects of varying physical conditions, e.g., temperature, turbidity plots were based on measurements taken after the completion of two periods: (i) an initial incubation period of protein, and nucleic acid, for 5 min, separately, i.e., prior to mixing, to allow the establishment of thermal and other (e.g., conformational) equilibria, and (ii) a subsequent incubation of 5 min after mixing of protein and DNA, to allow an LLPS equilibrium to be established, in addition to thermal and chemical equilibria. For turbidity plots of pH-dependence, and for microscopic imaging of the effect of mixing, proteins were subjected to buffer exchange in order to transfer them into various buffer systems using repeated washing and centrifugal-ultrafiltration in Merck centrifugal filter units (cat no. UFC500396). Solutions of different pH were created using citrate buffer (pH 4), Tris buffer (pH 6 and pH 8), and CAPS buffer (pH 10). Buffers of different pH all contained 150 mM KCl, by way of salt.

### Acidic Native PAGE

All proteins were loaded at a concentration of 1.2 mg/ml on acidic native gels (15 % acrylamide) that were cast and run according to the protocol mentioned at http://ubio.bioinfo.cnio.es/data/crystal/local_info/protocols_old/ANATPAGE.html. Gels were run at 60 V on ice in a Bio-Rad Mini-Protean Tetra vertical electrophoresis setup. Gels were run for approximately 3 hours and then stained using Coomassie brilliant blue staining solution.

### Fluorescence labeling

Cysteine mutants were labeled using Alexa Fluor™ 594 C_5_ Maleimide, excited by 594 nm light, or Alexa Fluor™ 488 C_5_ Maleimide, excited by 488 nm light, or Fluorescein-5-Maleimide, also excited by 488 nm light. 100 μM of protein was incubated with 300 μM of TECP (Tris(2-carboxyethyl) phosphine hydrochloride, from Merck) and 200 μM of dye. Free dye was removed by ultrafiltration using Merck centrifugal filter units (cat no. UFC500396). Labeling efficiencies were calculated according to the protocols of manufacturers using a scanning Cary 50 Bio UV-Vis spectrophotometer.

### Fluorescence Microscopy

Samples for microscopy were mixed on the bench and visualized immediately under either on a Leica widefield THUNDER imaging system, or on a Zeiss LSM 980 system, for all samples involving DNA, or RNA. In general, in all experiments in which protein (HU-B, HU-A, or Dps) was being visualized, the unlabeled form of the protein (99 %) was spiked with a fluorescently-labeled form of the protein (1 %). Makeshift chamber slides were made using double-sided tape, to prevent the crushing of droplets by coverslips. All images were analyzed using Fiji ImageJ. For experiments involving Poly(U) RNA, the Zeiss LSM 980 was used, and 500 nM of Alexa Fluor™ 594 C_5_ Maleimide labeled HU-B was spiked into the mixture, before the addition of RNA, for imaging of the labeled HU-B protein engaging in droplet formation, with associated turbidity, due to the addition of the Poly(U) RNA, as the RNA itself was not labeled for this experiment. Brightfield, DIC, and phase-contrast images were all collected on an Olympus IX-83 widefield fluorescence microscope.

### Super-resolution microscopy

Cells were aliquoted from *E. coli* cultures, in the log phase of growth, overexpressing RFP-HU-A, and then air-dried upon a slide for visualization using the 564 nm laser line of a super-resolution microscope (Zeiss Elyra 7) based on Lattice structured illumination microscopy (SIM) technology, with use of the Plan Apo 63×/1.40 oil objective, and an sCMOS camera (PCO Edge).

### Fluorescence recovery after photobleaching (FRAP) assays

FRAP assays were carried out on a Zeiss LSM 980 microscope. *<u>HU-B</u>*. HU-B (50 μM) was spiked with 1% of Alexa 488-labeled HU-B mutant (containing an N-terminal Cys conjugated with Alexa Fluor™ 488 C_5_ Maleimide from Thermofisher; Cat no. A10254). The 488 nm laser was set at 100% for efficient bleaching and focused on a small circular region with a diameter of 1 μm. Iterations of the incident laser were set at 150. Fluorescence intensities were measured for 100 cycles. Data was collected in triplicates and plotted for the same after normalization. *<u>HU-A</u>*. FRAP of HU-A was performed exactly as for HU-B, with the spiking of HU-A with 1 % labeled HU-A (i.e., the S17C mutant of HU-A labeled by the Alexa Fluor™ 488 C_5_ Maleimide fluorophore), with a different protein identity (HU-A), different concentration (55 μM) and different laser focus diameter (0.5 μm). *<u>Dps</u>*. FRAP of Dps was performed exactly as for HU-B, with the spiking of Dps with 1 % labeled Dps (i.e., the S22C mutant of Dps labeled by the Alexa Fluor™ 594 C_5_ Maleimide), with a different protein identity (Dps), different protein concentration (30 μM), different laser line (594 nm, instead of 488 nm) and the same laser focus diameter as used for HU-A (0.5 μm).

### Analytical size-exclusion chromatography to determine multimericity

Size exclusion chromatography (SEC) was performed on a Superdex-75 Increase column (GE), on an AKTA Purifier 10 workstation (GE), with equilibration of the column with Tris (50 mM) buffer of pH 7.4 containing 150 mM KCl, at room temperature. All protein loading was carried out using 500 μl protein of 110 μM concentration. Monitoring of the elution volume(s) of protein(s) was done through the measurement of the 280 nm UV absorption of the eluent.

### HU-induced DNA accretion experiment using Salmon Testis DNA

Salmon testis DNA (Stock concentration of 10 mg/ml; Sigma/Merck Cat no. D-9156) was labeled with SYBR Green dye (Stock concentrations 10000X) by adding dye (final concentration of 1 μl/10 ml in solution with DNA) to DNA (~100 μl), following which free SYBR Green dye was removed through repeated washing and centrifugal-ultrafiltration in Merck centrifugal filter units (cat no. UFC500396). The resulting labeled DNA (20 μl volume; concentration not determined) was pipetted onto a cover slip and visualized on an inverted Nikon Eclipse Ti-u microscope, as a uniform field of green fluorescence. Following this, with continuing video-based visualization of this drop of 20 μl, using the microscope’s camera, 20 μl of HU-B (100 μM) protein was added to the DNA already present (and being visualized) on the coverslip, and the mixed drop containing HU-B and Salmon testis DNA was visualized over time.

### Examination of DNA polymerase’s ability to associate with DNA in HU-B-condensates

Pfu DNA polymerase (purified from a bacterial clone expressing the polymerase previously produced in our laboratory) was labeled, using Alexa Fluor™ 594 C_5_ Maleimide. The labeled DNA polymerase was then added to Venus-HU-B using standard recommended buffer conditions, and droplets formed after the addition of 4WJ DNA (with associated solution turbidification) were imaged using confocal microscopy, as for other experiments, to examine whether the labeled DNA polymerase colocalized with the HU-B and the DNA, within droplets.

## Acknowledgments

We thank the Government of India for financial support (for equipment/consumables) extended through IISER Mohali, the Ministry of Education (Centre of Excellence grant in Protein Science, Design and Engineering to S.M. and P.G.), the Department of Biotechnology (HEHRC grant to P.G.), and the Department of Science and Technology (Nano-Mission grant to S.M. & FIST grant to the Department of Biological Sciences, IISER Mohali). A.G., A.J, and K.A., thank DBT (India), IISER Mohali and UGC (India), respectively, for research fellowships.

## Authors contributions

All authors participated in the work’s conception, and discussion of results. Experiments were performed by A.G., along with A.J. (HU-B phase separation) or K.A. (bacterial imaging). A.G. & P.G. prepared figures, and text drafts, with seminal contributions from A.J. & S.M. Overall directions, supervision and editing came from S.M. & P.G.

## Competing interests

The authors declare no conflict of interests.

## Data availability

The data are available within the Article, Supplementary Information, and Source Data file.

## Notes

### Competing Interest Statement

The authors have declared no competing interest.

