## Supplementary Information for "Nucleoid-Associated Proteins Undergo Liquid-Liquid Phase Separation with DNA into Multiphasic Condensates Resembling Bacterial Nucleoids"

[§]Contributed equally

**a**

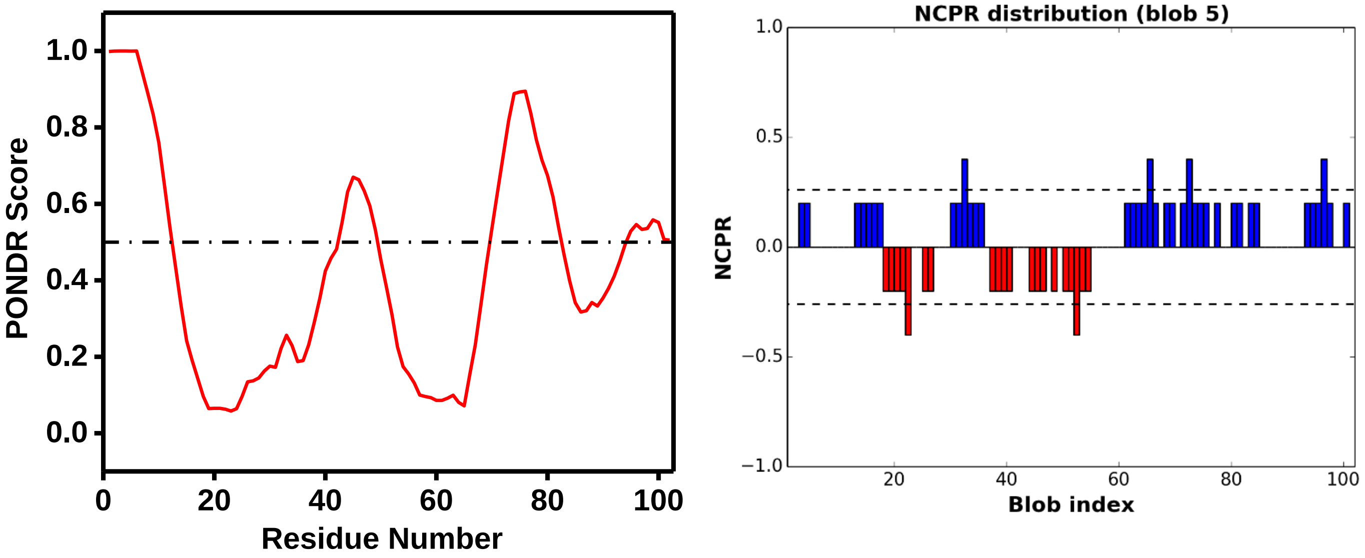


**b**

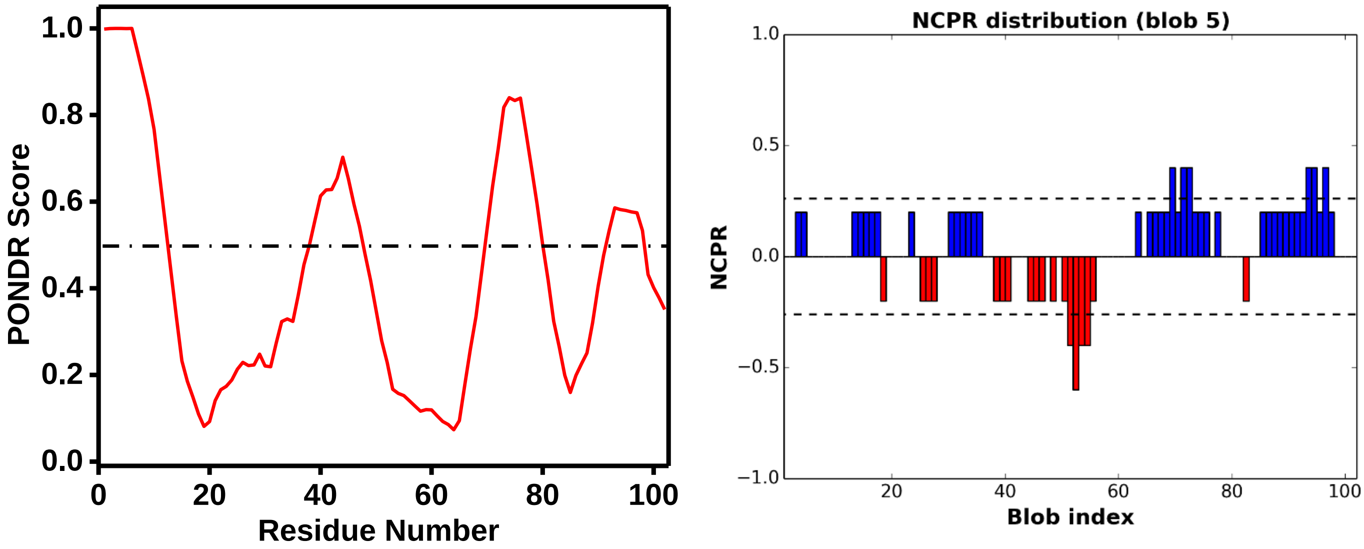


**Supplementary Figure 1:** PONDR and CIDER plots showing significant amounts of disorder and charge clusters in **a.** HU-A, **b** HU-B.

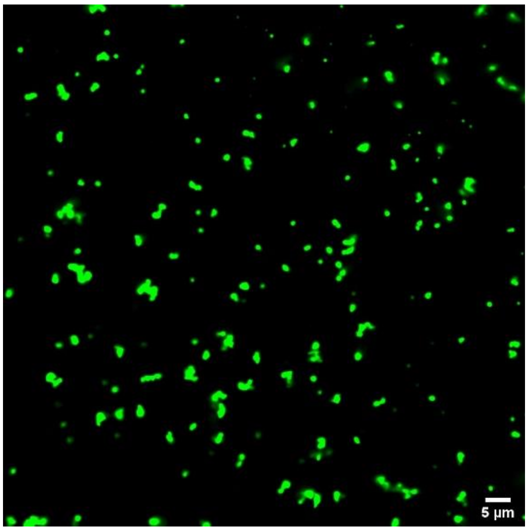


**Supplementary Figure 2:** Confocal microscopic image of the condensates shown in Supplementary Video 2.

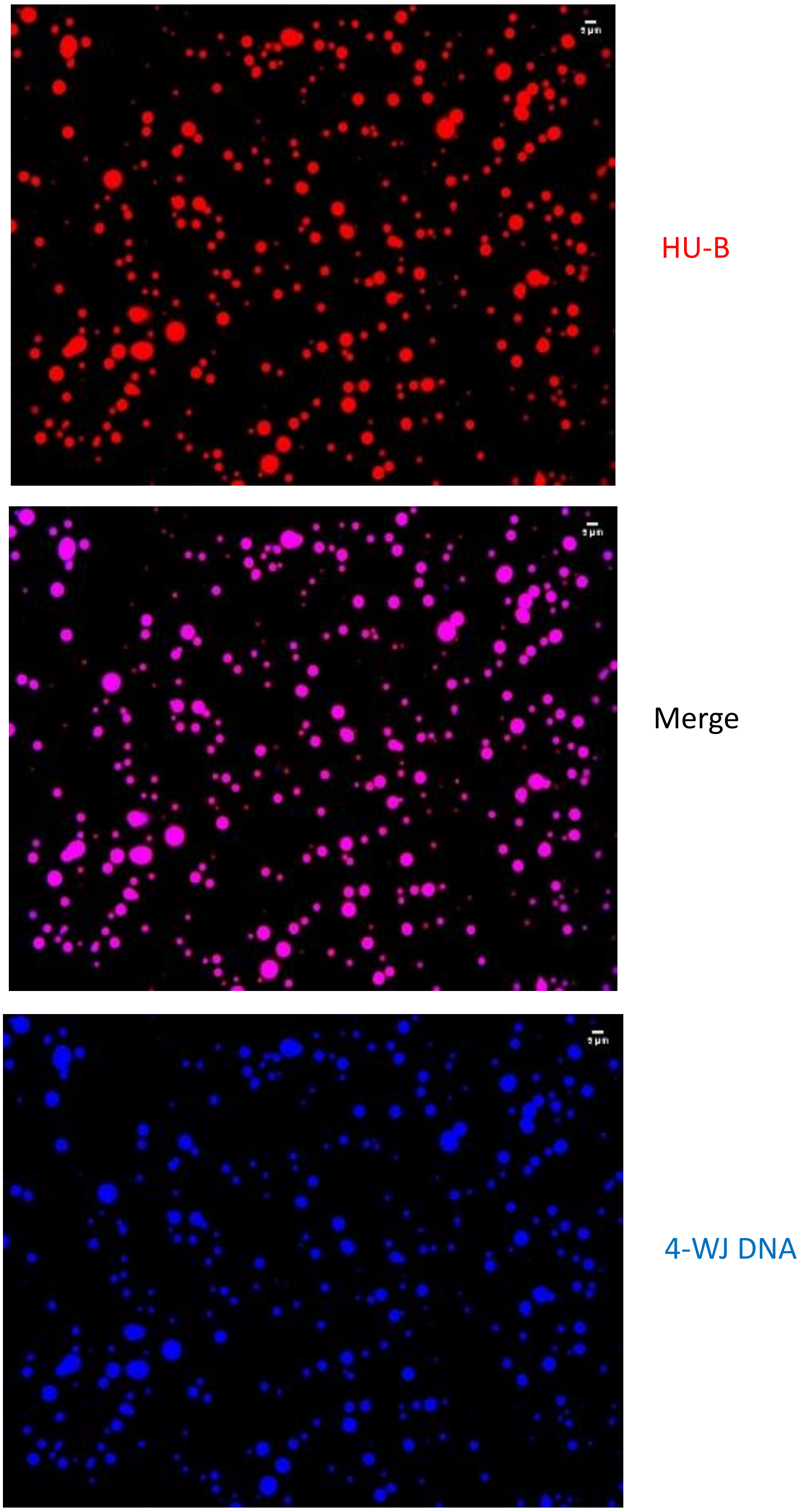


**Supplementary Figure 3:** Phase separation of HU-B into liquid condensates in the absence of PEG 6000, in the presence of 5 µM 4WJ DNA (rather than 3 µM 4WJ DNA).

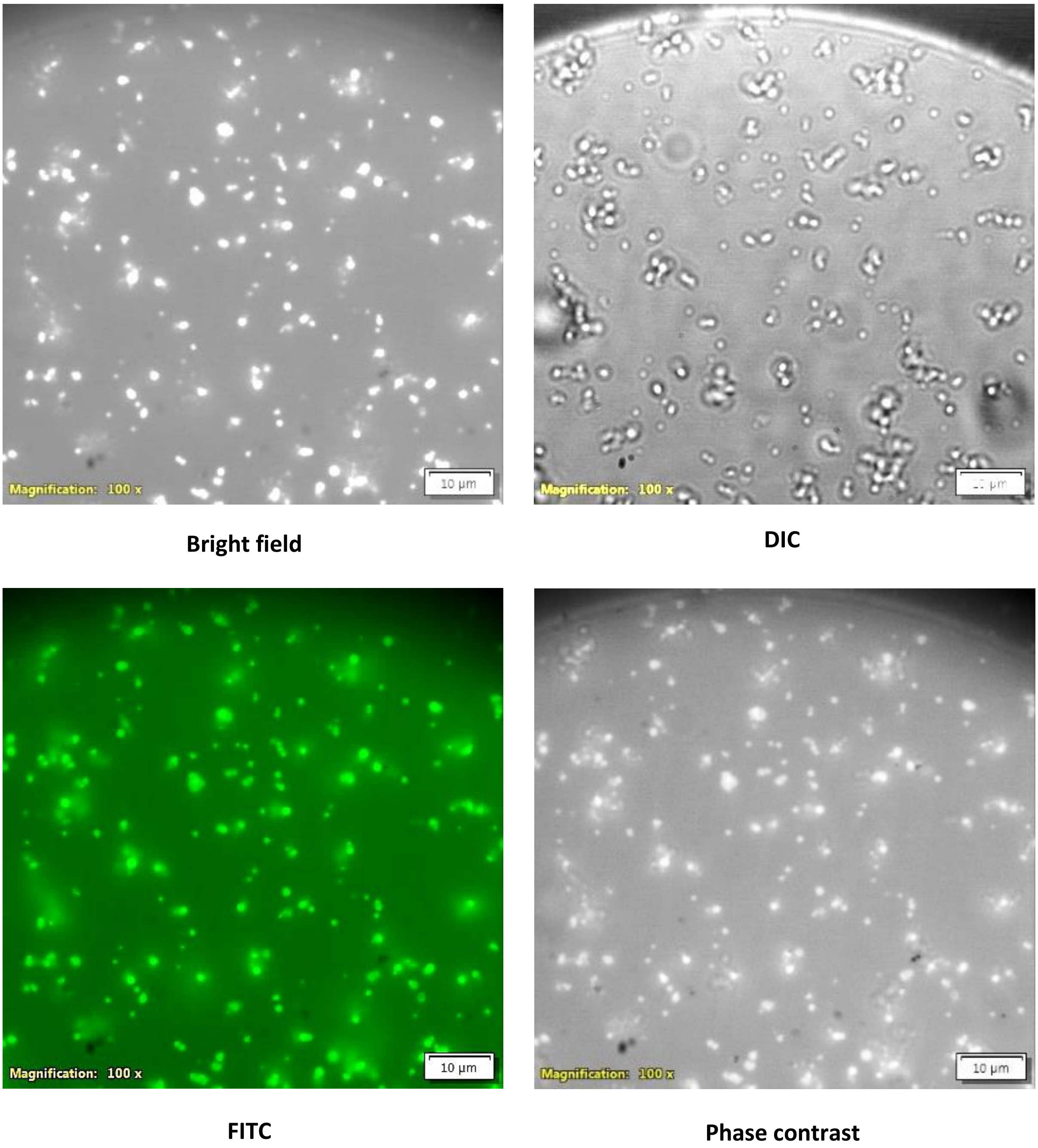


**Supplementary Figure 4:** Phase-separated HU-B condensates visualized using bright field, DIC, fluorescence (FITC), and phase-contrast microscopic techniques.

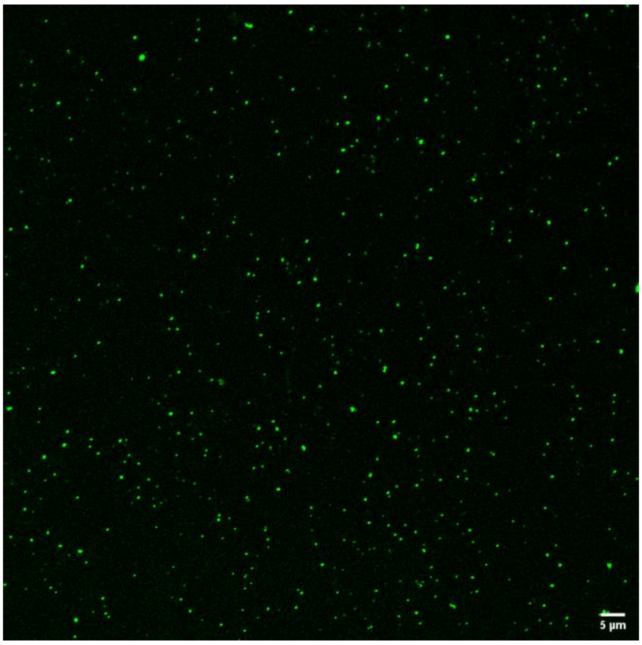


**Supplementary Figure 5:** Phase-separated HU-B condensates formed in absence of DNA, in the presence of 8 % PEG 6000 (rather than 2 % PEG 6000.

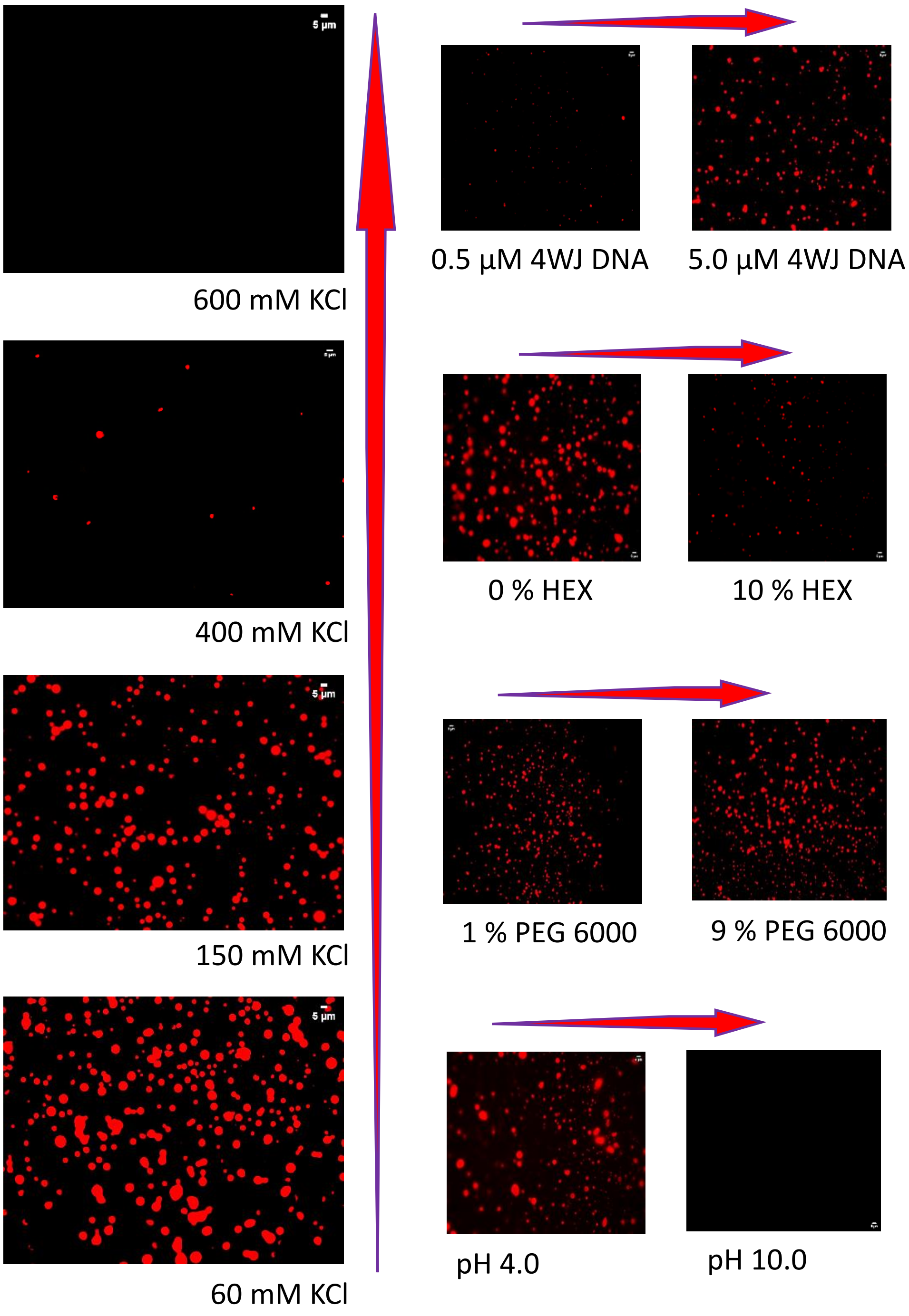


**Supplementary Figure 6:** Microscopic images corresponding to the turbidity plots shown in Figure 2 in the main manuscript.

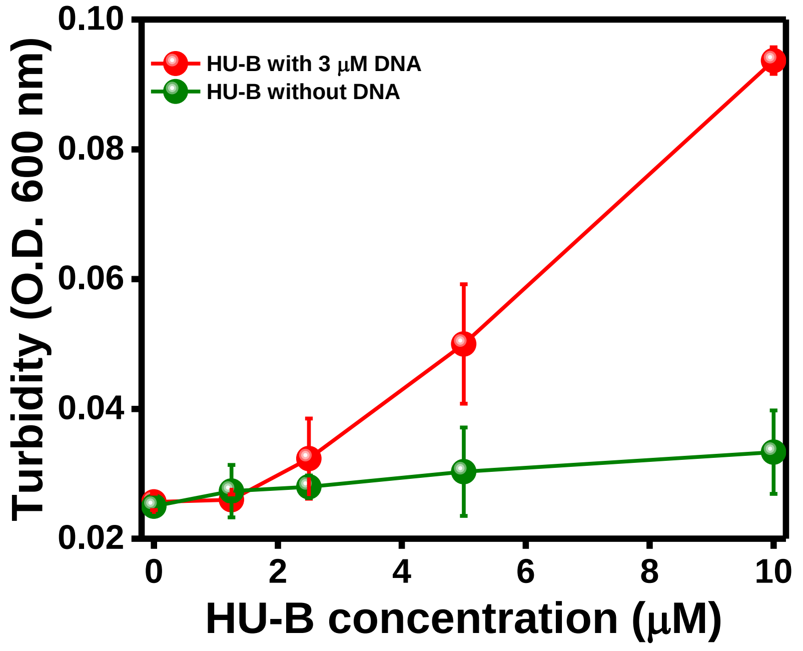


**Supplementary Figure 7:** Enlarged section of Figure 2**b,** highlighting the $c_{sat}$ value of HU-B, under the standardized conditions, to be approximately 2.5 μM.

a

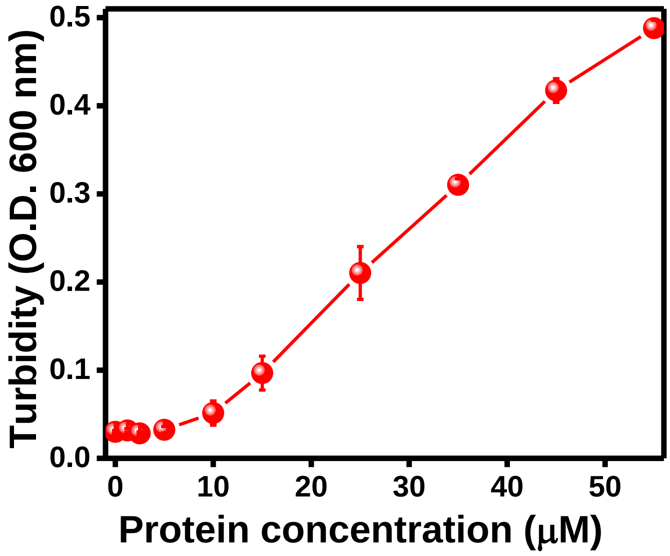


b

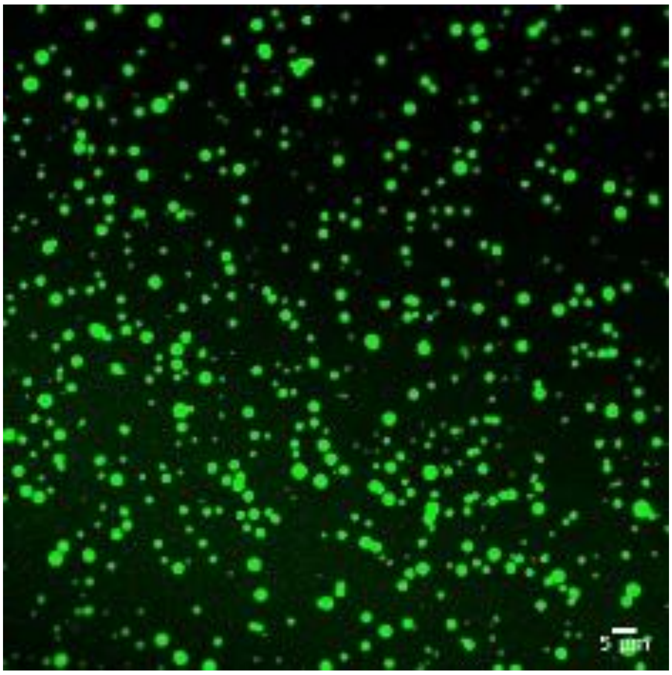


**Supplementary Figure 8:** Behaviour of the HU-BA simulacrum. **a** Turbidity plot, showing phase-separation by the HU-BA simulacrum at low protein concentrations, with a $c_{sat}$ value of 5-10 μM. **b** Representative confocal microscopic image showing condensates formed by the HU–BA simulacrum.

**a**

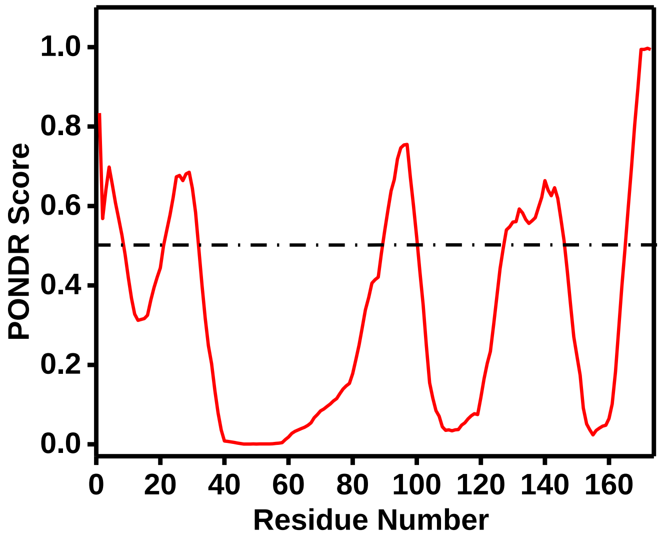


**b**

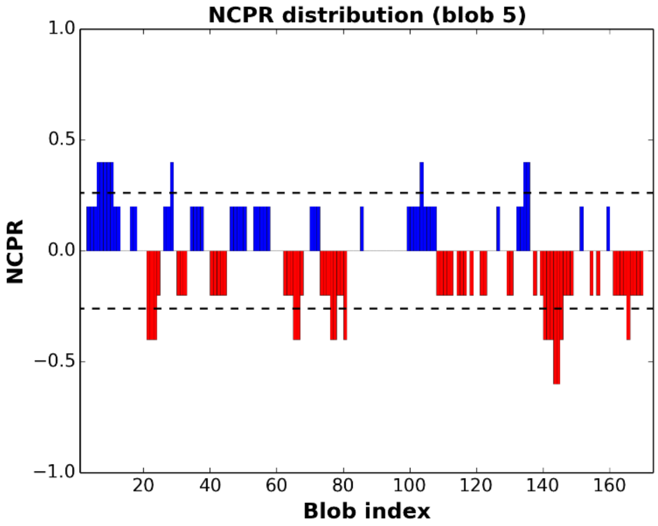


**c**

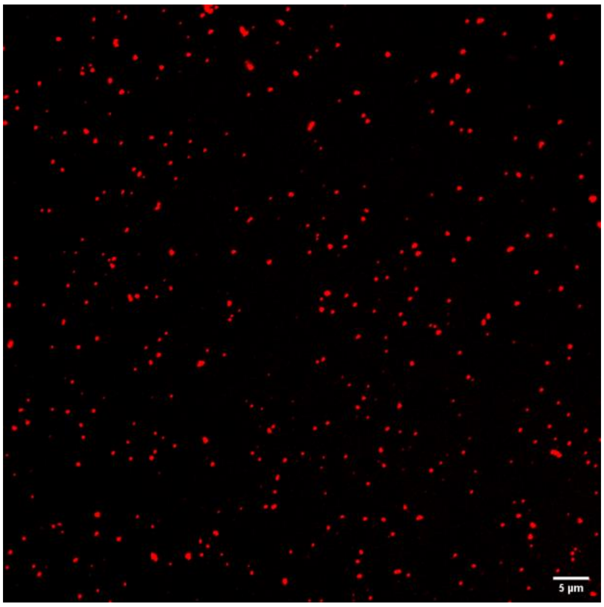


**Supplementary Figure 9:** PONDR and CIDER **(a,b)** plots showing significant amounts of disorder and charge clusters respectively in Dps. **(c)** Formation of Dps condensates in absence of DNA, in the presence of 8 % PEG 6000.

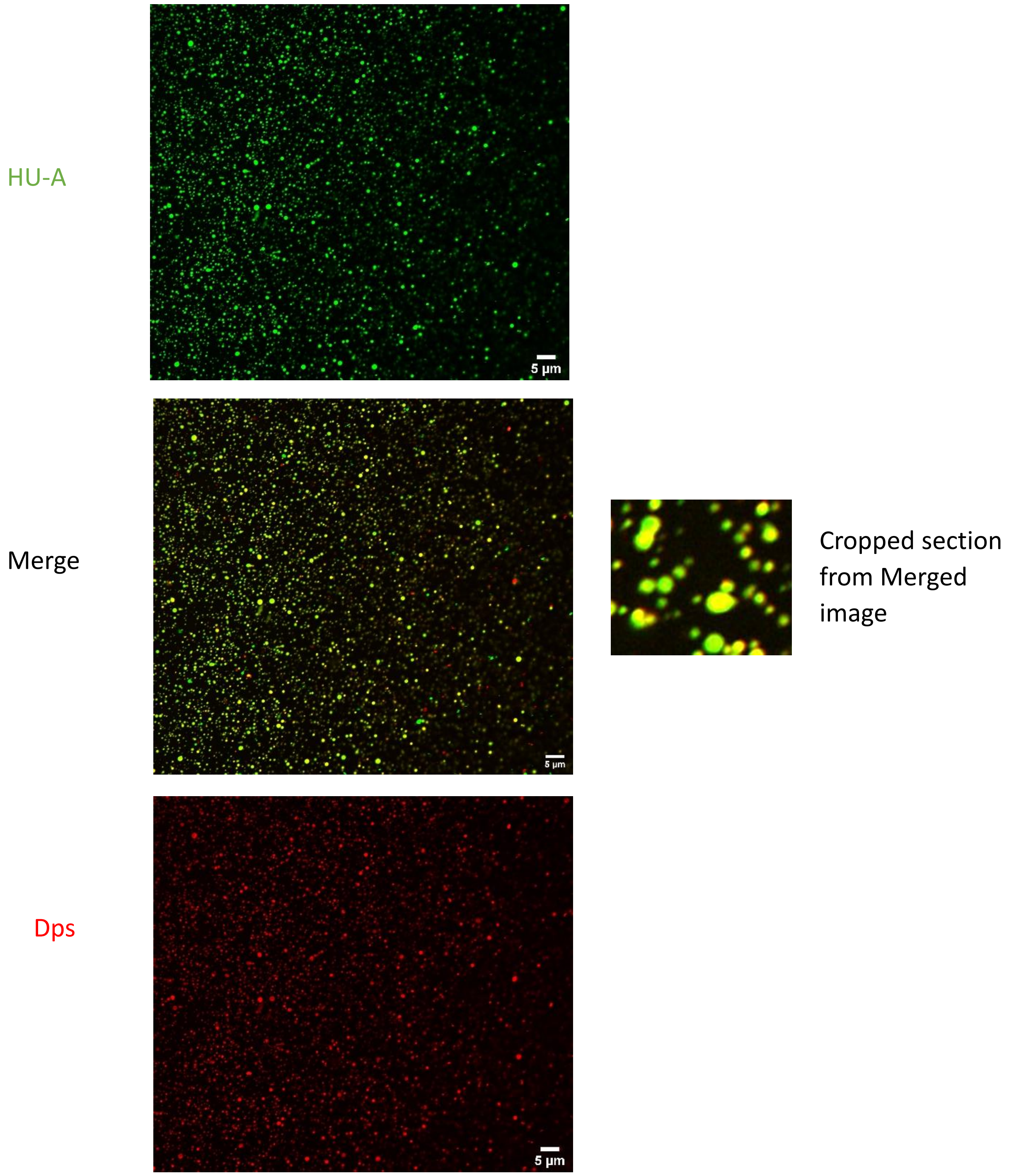


**Supplementary Figure 10:** Droplet in droplet behaviour seen also in coacervation of HU-A and Dps with 4WJ DNA (as shown for HU-B and Dps with 4WJ DNA in Figure 5 in the main manuscript).

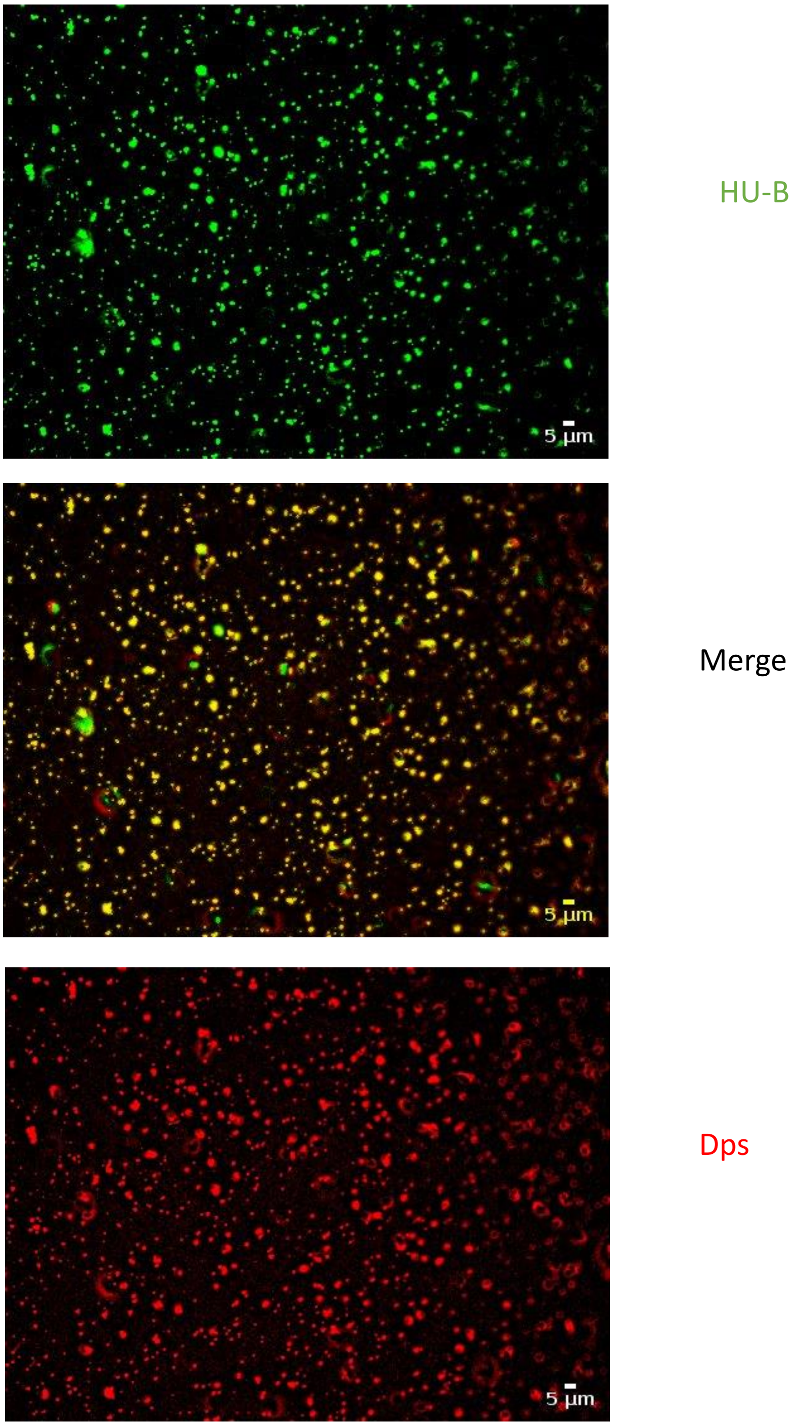


**Supplementary Figure 11:** Coacervation of HU-B and Dps in the absence of 4WJ DNA, in the presence of 8% PEG 6000. No droplet-in-droplet structures are visible here.

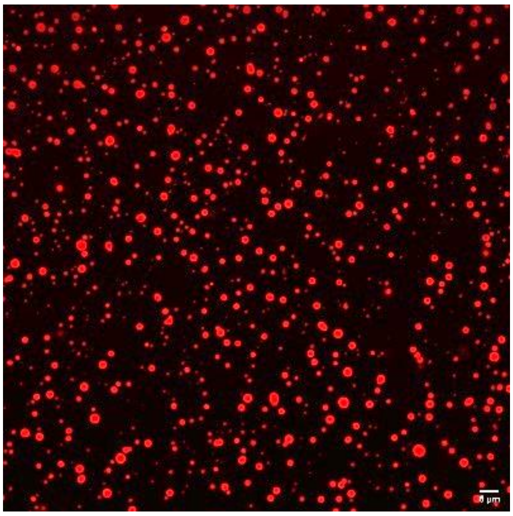

**Supplementary Figure 12:** Condensates of HU-B created using 160 ng/ul Poly(U) RNA, instead of DNA.

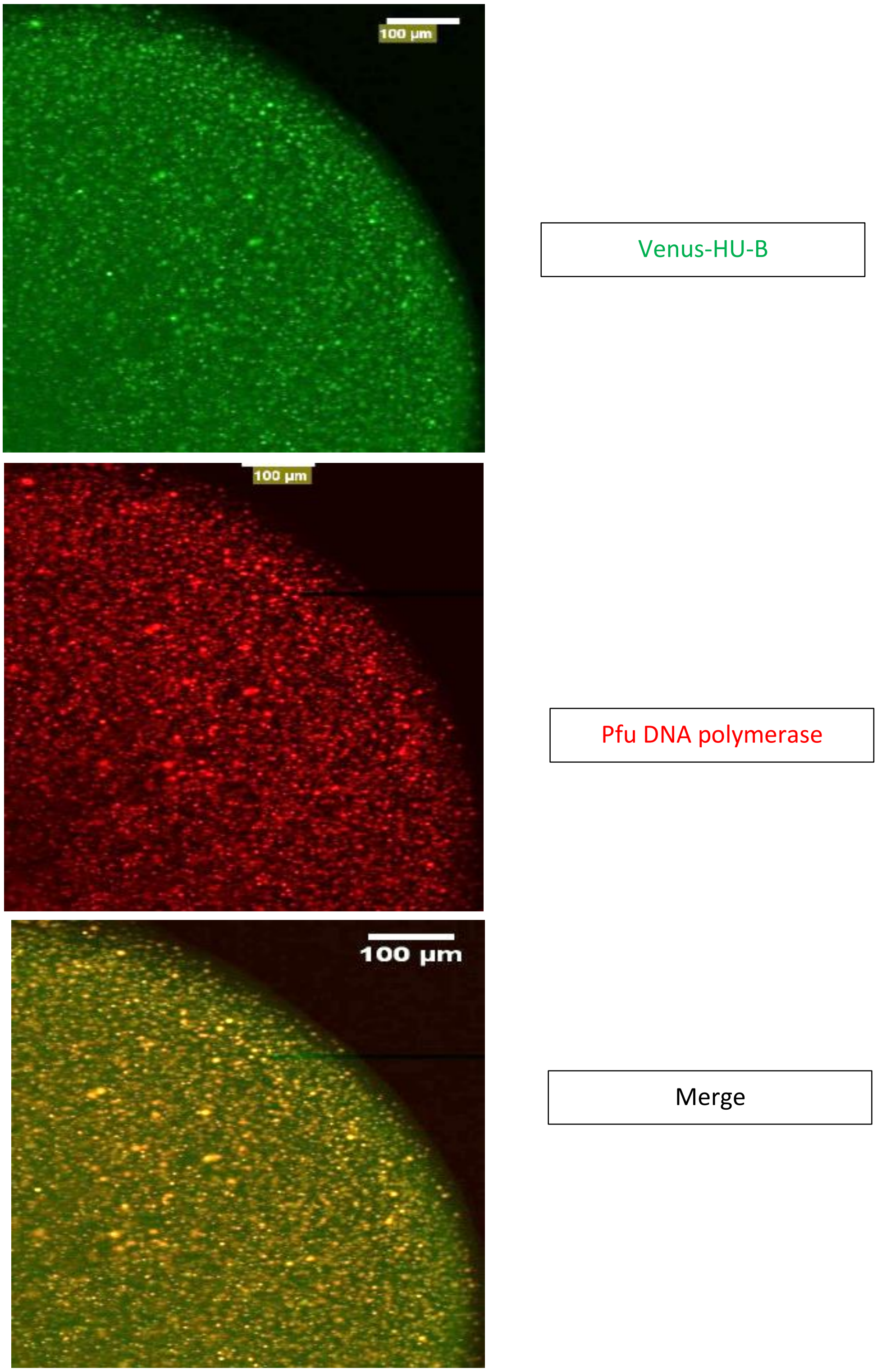


**Supplementary Figure 13:** Colocalization of Pfu DNA polymerase with Venus-HU-B.
